# Role of ADMA-histones in dual-strand piRNA source loci recognition by Rhino

**DOI:** 10.1101/2024.03.15.585151

**Authors:** Raku Saito, Hirotsugu Ishizu, Ritsuko Harigai, Kensaku Murano, Yurika Namba, Mikiko C. Siomi

## Abstract

In *Drosophila* germ cells, piRNAs arise from dual-strand piRNA clusters, which are marked by a repressive histone mark H3K9me3, but are transcribed from internal sites in a manner dependent on the HP1 homolog Rhino. Rhino binds to H3K9me3 on these clusters, yet the mechanism controlling its binding to clusters remains unclear. Here, we used cultured ovarian somatic cells (OSCs), which lack endogenous Rhino and its stabilizer Kipferl, whose absence renders the dual-strand clusters inert, and found that exogenous Rhino tends to bind to the ends of dual-strand clusters with asymmetric dimethylarginine histones (ADMA-histones). Depletion of the arginine methyltransferases responsible for ADMA modifications, such as DART4, affected the genomic localization of Rhino in both OSCs and ovaries. We also identified primitive, cluster-like genomic regions, termed DART4 piSL, where Rhino propagates in an DART4-dependent but unstable manner. Our study proposes that ADMA-histones play a crucial role in the initial genome loading of Rhino and may establish the potential sites of its propagation, which are subsequently stabilized to support piRNA production.

## Introduction

PIWI-interacting RNAs (piRNAs) are germline-enriched small non-coding RNAs that repress transposons in collaboration with PIWI proteins to maintain fertility^1,2,3^. In *Drosophila* germ cells, piRNAs are primarily derived from dual-strand piRNA clusters, intergenic elements rich in transposon fragments^4,5^. These clusters are occupied with the repressive histone mark H3K9me3 and lack typical promoters, but are transcribed in both directions from the internal sites^6,7^. This unconventional transcription requires special factors, including heterochromatin protein 1 (HP1) homolog HP1d/Rhino (Rhino), and provides piRNA precursors with a variety of sequences, which are then processed to mature piRNAs in the cytoplasm^5,6,7,8,9,10,11,12,13,14^. The mechanism dedicated to this piRNA processing is known as the ping-pong cycle, where Aubergine and Argonaute 3 cleave the precursors in a manner dependent on the sequences of piRNAs already bound to them^1,2,3^. This pathway also consumes transposon RNA transcripts as the sources of mature piRNAs, resulting in transposon repression at the post-transcriptional level^2,3^.

Like other HP1 family members, Rhino has a chromodomain through which it binds to H3K9me3 scattered on dual-strand piRNA clusters^6,8,9,12,13,14,15,16^. Deadlock and Cutoff then join Rhino, assembling the Rhino-Deadlock-Cutoff (RDC) complex, which further recruits the transcription factor IIA-L (TFIIA-L) paralogue Moonshiner, the transcription factor IIA-S (TFIIA-S), and the animal TATA-box-binding protein (TBP) paralog TRF2, initiating transcription of the clusters^6,7,10^. Upon transcription, Cutoff binds to piRNA precursors and protects them from splicing, contributing to the production of an abundance of piRNAs^17,18,19,20^.

A recent study identified the zinc-finger protein Kipferl as a factor required for Rhino to distinguish between H3K9me3 located on transposons and dual-strand piRNA clusters^12^. Kipferl stabilizes Rhino on guanine-rich sequence motifs present in dual-strand clusters in an H3K9me3-independent manner^12^. Maternally inherited piRNAs also play a role in the localization of Rhino to dual-strand clusters during early embryogenesis via Piwi-dependent H3K9me3 epigenetic marks ^9,21^.

Rhino has a propensity to spread across the genome. For example, depletion of the histone demethylase Kdm3 in the female germline causes Rhino to spread to adjacent chromatin^13^. Deletion of the region proximal to the transcription start site of the *phospholipase D* gene, which is in the region flanking piRNA cluster 42AB, also resulted in Rhino spreading toward the deleted genomic region^10^. In addition, Rhino tethering experiments with artificial constructs showed that Rhino spreads as the constructs are transcribed^7^. In each case, production of piRNAs from regions after Rhino spreading was detected. These findings suggest that piRNA clusters are defined by Rhino spreading over the genome and imply that the process of piRNA cluster formation is composed of three steps: Rhino spreading, Kipferl-dependent stabilization of Rhino-genome association, and epigenetic inheritance of the Rhino positional memory by maternally inherited piRNAs. Since the genomic localization of both Rhino and Kipferl is interdependent, it is hypothesized that the initial loading of Rhino on the genome precedes its spread or stabilization, and that this process is independent of Kipferl. To investigate this, it is necessary to observe the behavior of Rhino in a cell environment where piRNAs are functional, but where both Kipferl and Rhino are not expressed. However, the mechanism of such Rhino behavior remains unclear.

In this study, we describe the important role of <u>a</u>symmetric <u>dim</u>ethyl<u>a</u>rginine histones (<u>ADMA</u>-histones) in the initial loading of Rhino onto the genome and their stabilization. We first established cultured ovarian somatic cells (OSCs)^22^ expressing FLAG-Rhino by induction with doxycycline (Dox). Unlike ovarian germ cells, OSCs lack Rhino and other factors required for the activation of dual-strand piRNA clusters, such as Deadlock and Cutoff^23^; therefore, dual-strand clusters are not in an active state for piRNA biogenesis^24^. We found that Rhino in OSCs assembles nuclear foci similar to Rhino foci in the ovary. We then performed chromatin immunoprecipitation sequencing (ChIP-seq) and showed that Rhino tends to bind to the ends of dual-strand clusters with ADMA-histones, such as H3R17me2a. Depletion of the arginine methyltransferases responsible for ADMA modifications, such as *Drosophila* arginine methyltransferase 4 (DART4), affected the genomic localization of Rhino in OSCs and in the ovary. We also identified genomic regions that have ADMA-histones at their ends and exhibit broad Rhino spreading across their internal regions in a DART4-dependent manner. In contrast to authentic piRNA clusters, Kipferl was lost together with Rhino upon DART4 depletion in these regions, suggesting that Kipferl by itself is not sufficient to stabilize Rhino binding; rather, their localization depends on DART4. piRNAs were mapped to these regions, albeit at low levels. The levels of these piRNAs decreased also in a DART4-dependent manner. Therefore, these regions, referred to as DART4-dependent piRNA source loci (DART4 piSL), might be nascent dual-strand piRNA source loci driven by the propagation of Rhino initially loaded onto the genome via ADMA-histones.

## Results

### Rhino localizes to the ends of dual-strand clusters in the OSC genome

To observe how Rhino in OSCs behaves intracellularly, we first performed immunofluorescence in FLAG-Rhino inducible OSCs to examine the subcellular localization of FLAG-Rhino upon Dox induction. This showed that Rhino assembled nuclear speckles in OSCs similar to Rhino-positive foci in the ovary^5,10^ when the cells were treated with Dox at a concentration of 5 ng/mL (Fig. 1A). At lower Dox concentrations and lower levels of Rhino expression, no clear foci appeared (Supplementary Fig. 1A, B), suggesting that Rhino-positive foci form in a manner dependent on the level of Rhino expression.

**Figure 1.**
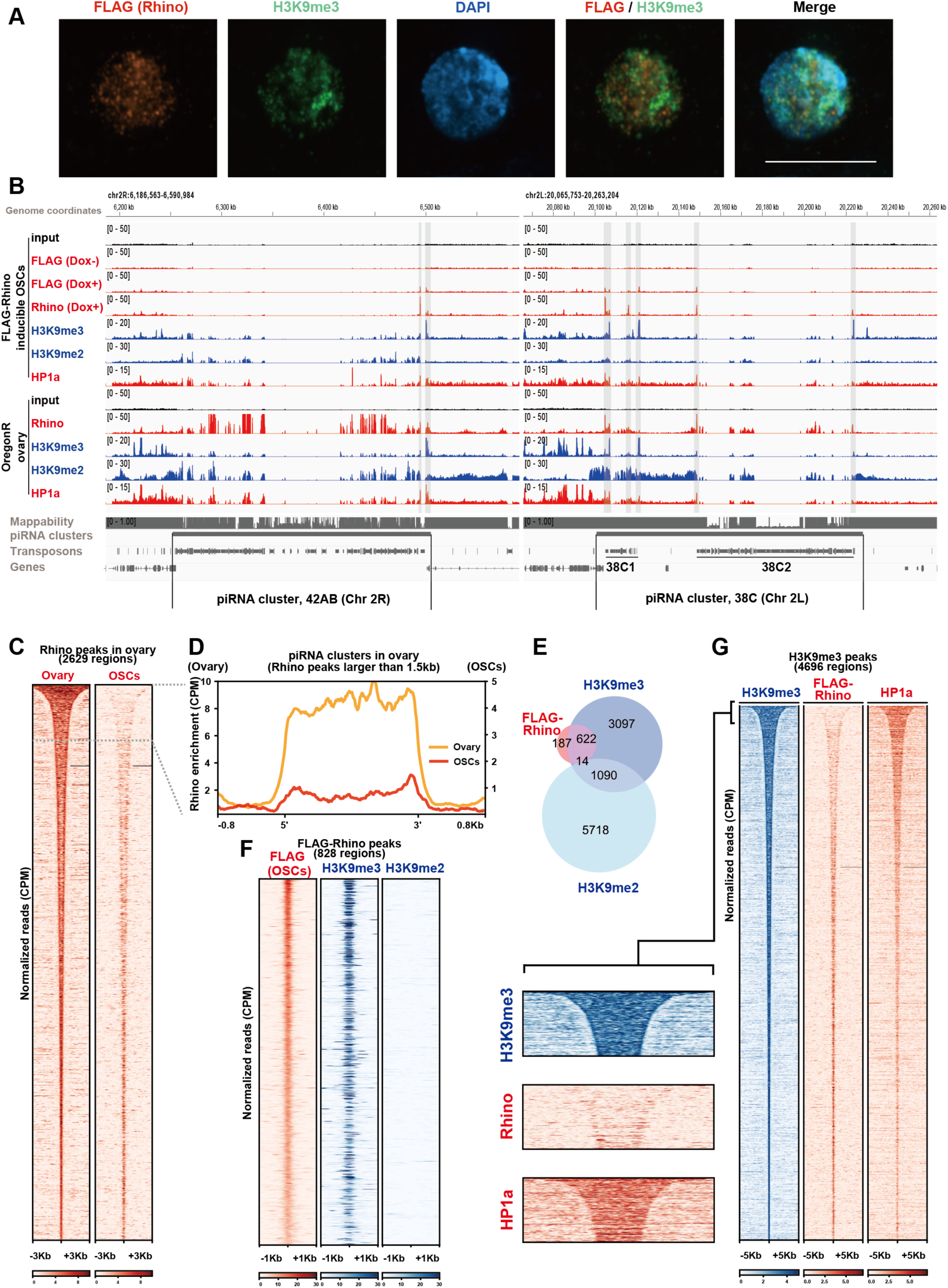
Rhino forms nuclear foci and binds the genome in OSCs. (A) Confocal immunofluorescent images of FLAG-Rhino (red) in FLAG-Rhino inducible OSCs upon Dox induction (5 ng/mL). H3K9me3 and OSC nucleus (DAPI staining) are shown in green and blue. Scale bar represents 10 µm. (B) Genome snapshot showing ChIP-seq in major dual-strand piRNA clusters (42AB and 38C). The gray-shaded areas indicate the sites at which FLAG-Rhino is localized in each piRNA cluster in OSCs. piRNA cluster 38C has two piRNA-producing regions, labeled 38C1 and 38C2. (C) Heatmap showing FLAG-Rhino localization of OSCs in Rhino peaks of *Drosophila* wild-type Oregon-R ovary. The regions of 6 kb around the Rhino localization site were plotted based on macs2 callpeak results. (D) Metaplot showing the localization of FLAG-Rhino in OSCs at piRNA clusters. For piRNA clusters, broad peaks of Rhino localization in the ovary greater than 1.5 kb were used. The overlap of each Rhino peak with annotated piRNA clusters (e.g., 42AB, 38C, and 80F) was identified, as listed in Supplementary Table 9. (E) Venn diagram showing the overlap of Rhino, H3K9me3, and H3K9me2 peaks in OSCs. (F) Heatmap showing the signals of ChIP-seq using anti-FLAG, anti-H3K9me3, and anti-H3K9me2 in FLAG-Rhino peaks of OSCs. The regions of 2 kb around the FLAG-Rhino localization sites were plotted based on macs2 callpeak results. (G) Heatmap showing the signals of ChIP-seq using anti-H3K9me3, anti-FLAG, anti-HP1a antibodies at H3K9me3 broad peaks of OSCs. The regions of 10 kb around the FLAG-Rhino localization sites were plotted based on macs2 callpeak results.

We next conducted ChIP-seq of FLAG-Rhino under conditions where Rhino assembles nuclear foci as in Fig. 1A. Genome mapping of the ChIP-seq reads showed that Rhino can be found, albeit weak, on dual-strand piRNA clusters, such as 42AB and 38C on chromosome 2, 80F on chromosome 3, and 102F on chromosome 4, all of which are in an active state in piRNA production in the ovary, but the signals seem to be biased toward the ends of the clusters (Fig. 1B and Supplementary Fig. 1C). ChIP-seq of Rhino was also performed in the ovary: Rhino was strongly enriched at highly transcribed piRNA clusters, 42AB, 38C, 80F, and 102F, as previously reported^5,6,7,8,9,10,11,12,13,14^ (Fig. 1B and Supplementary Fig. 1C), which suggests that our ChIP-seq procedure is reliable.

We then compared the mode of Rhino genome binding in OSCs with that in the ovary in a more comprehensive manner. The results show that Rhino expressed in OSCs bound predominantly to genomic sites exhibiting sharp and interspersed Rhino localization patterns in the ovary, while showing little localization within broad Rhino domains, including major piRNA cluster (Fig. 1C). The metaplot indicates a tendency for Rhino in OSCs to be toward the terminal regions of broad domains, such as the cluster ends (Fig. 1D). Rhino also binds to the cluster bodies in OSCs, but the signals were weaker compared to those at the ends. Based on these results, we interpret that OSCs contain a factor(s) that promotes Rhino binding toward the ends of the clusters, but unlike the ovary, they lack a factor(s) that promotes and stabilizes Rhino spreading throughout the clusters.

The terminal localization of Rhino in OSCs does not appear to be due to H3K9me2 (Fig. 1B, E, F): Of the 823 Rhino signals, only 14 signals overlapped with the H3K9me2 signals (Fig. 1E). The heatmap also showed that H3K9me2 barely accumulated at the Rhino location (Fig. 1F). The positional correlation between Rhino and H3K9me3 is stronger than between Rhino and H3K9me2 (i.e., 77%; 636 in 823 signals) (Fig. 1E). Nonetheless, the overlap between Rhino and H3K9me3 was not very strong according to the heatmap (Fig. 1F, G). Immunofluorescence also showed weak overlap between the prominent H3K9me3 speckles and FLAG-Rhino foci (Fig. 1A). Since immunofluorescence primarily visualizes H3K9me3 foci that correspond to broad heterochromatic domains in the genome, such as those at centromeres, pericentromeres, or telomeres (named chromocenters), the sharp and interspersed H3K9me3 signals along chromosome arms, where Rhino partially colocalizes (Fig. 1F), are difficult to visualize, resulting in little apparent overlap with Rhino (Fig. 1A). HP1a, another HP1 family member required for constitutive heterochromatin and known to be a general H3K9me3 marker, co-localizes with H3K9me2/3 and is distributed along the entire length of piRNA clusters similarly in both OSCs and ovary (Fig. 1B and Supplementary Fig. 1C). HP1a is present in the broad H3K9me3 peak (Fig. 1G). Therefore, although H3K9me3 and HP1a might be involved in some aspect of the mechanism underlying Rhino localization, they do not contribute to the localization specificity within that mechanism at least.

FLAG-Rhino ChIP-seq was also performed at low Rhino expression levels. The trend toward terminal localization of Rhino at 5 ng/mL Dox was observed at 0.05 ng/mL and 0.5 ng/mL Dox as well (Supplementary Fig. 1D, E). This indicates that Rhino binds to the OSC genome even at low expression levels. However, Rhino foci were not clearly observed with the lower Dox concentrations (Supplementary Fig. 1A). It is interpreted that genome binding occurs first, followed by Rhino foci formation, which is dependent on Rhino expression levels.

The use of anti-FLAG antibodies in Rhino ChIP in OSCs was justified by the high similarity between the signals of anti-FLAG and anti-Rhino antibodies (Fig. 1B and Supplementary Fig. 1C, D).

### Rhino co-localizes with asymmetric dimethylarginine histones on the OSC genome

To identify genomic scaffolds that are potentially involved in determining the genomic localization of Rhino in OSCs, we performed ChIP-seq of key histone modifications and compared the data with Rhino ChIP-seq data. Heatmap revealed that specific ADMA-histones, including H3R2me2a, H3R8me2a, H3R17me2a, and H4R3me2a, tend to co-localize with Rhino across the OSC genome (Fig. 2A). Consistent with this, these ADMA-histones showed a tendency to localize to the ends of Rhino-bound dual-strand piRNA clusters (Fig. 2B, C, and Supplementary Fig. 2A). H3K4me1 and H3K27ac have been reported to co-localize with H3R8me2a in mammals^25^, which was not observed in OSCs (Fig. 2A, B, and Supplementary Fig. 2A). Co-localization of H3K4me1 and H3K27ac with Rhino was also not observed.

**Figure 2.**
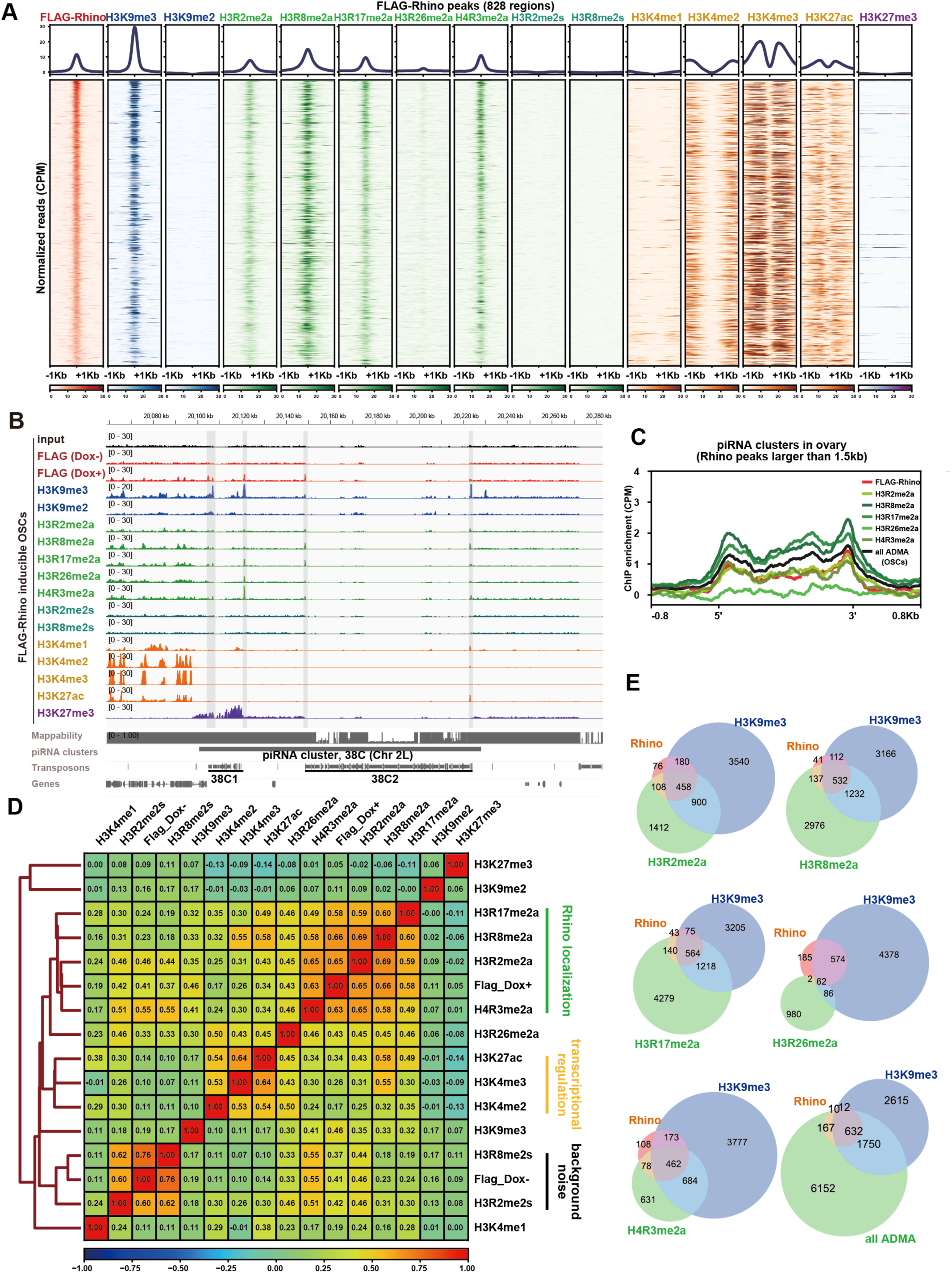
ADMA-histones tend to co-localize with FLAG-Rhino in OSCs. (A) Heatmap showing key histone modifications and HP1a ChIP-seq results at FLAG-Rhino localization sites (828 regions) in OSCs. The regions of 2 kb around the FLAG-Rhino localization sites were plotted based on macs2 callpeak results. (B) Genome snapshot showing all histone modifications’ ChIP-seq data in major dual-strand piRNA clusters (38C). The gray-shaded areas indicate the localization sites of FLAG-Rhino in each piRNA cluster in OSCs. (C) Metaplot showing the localization of ADMA-histones and FLAG-Rhino in OSCs at piRNA clusters. For piRNA clusters, broad peaks of Rhino localization in the ovary greater than 1.5 kb were used. (D) Pearson correlation coefficient comparing the coverage per 100 bp bin size of all ChIP-seq mapping results to the dm6 genome and its hierarchical clustering. (E) Venn diagram showing sites of overlapping localization of FLAG-Rhino, H3K9me3, H3R2me2a, H3R8me2a, H3R17me2a, H3R26me2a, H4R3me2a, and all ADMA-histones in OSCs.

Examination of the correlation coefficients revealed that signal peaks of each ADMA-histone were significantly correlated with those of FLAG-Rhino (Fig. 2D). The ChIP-seq signals of H3R2me2s and H3R8me2s correlate more strongly with FLAG(Dox-) than with FLAG(Dox+), suggesting that these histone modifications may represent background noise (see “background noise” in Fig. 2D). In contrast, ADMA-histones show a higher correlation coefficient with FLAG(Dox+) than with FLAG(Dox-): it is also worth noting that in hierarchical clustering, ADMA-histones form a monophyletic group with FLAG(Dox+) (see “Rhino localization” in Fig. 2D). Histone modifications related to transcription, such as H3K4 and H3K27, as well as constitutive heterochromatin markers like H3K9me2 are not associated with the localization of FLAG-Rhino (see “transcriptional regulation” in Fig. 2D). Correlation of ADMA-histones in the genomic localization of Rhino was implicated.

The overlaps of ChIP-seq peaks were examined. In the diagram shown in Fig. 1E, approximately 23% of Rhino ChIP peaks did not overlap with H3K9me3 (187 in 823 sites), but these Rhino peaks were almost encapsulated by the read sets of ADMA-histones (Fig. 2E). This suggests that ADMA-histones are involved in the localization of Rhino. However, the degree of overlap of those sites is not very high in each case (Supplementary Fig. 2B). It appears that each ADMA-histone contributes differently to Rhino localization. HP1a was found at the dual-strand clusters in OSCs (Fig. 1B). However, unlike Rhino, genomic localization of HP1a does not appear to be determined by ADMA-histones and H3K9me3 because approximately 30% of HP1a ChIP peaks did not overlap with the read sets of ADMA-histones and H3K9me3 (492 in 1,672 sites) (Supplementary Fig. 2C). These results raise the possibility that ADMA-histones, together with H3K9me3, may contribute specifically to the recruitment of Rhino to the ends of dual-strand clusters in OSCs.

### Arginine methyltransferases DART1 and DART4 differently regulate Rhino genomic binding

Arginine methylation of histones is catalyzed by protein arginine methyltransferases (PRMTs), and the *Drosophila* genome encodes 11 of these PRMTs, also known as *Drosophila* arginine methyltransferases (DARTs)^26,27^. Of those, DART1, a homolog of mammalian PRMT1, and DART4/CARMER (DART4), a homolog of mammalian PRMT4/CARM1 (CARM1), were reported to be type I PRMTs that catalyze the methylation of monomethylarginine to asymmetric dimethylarginine^28,29,30,31,32,33,34,35^. DART8 is involved in asymmetric dimethylation of H3R2^36^. CG17726 is considered to be a DART member because of its sequence similarity and is expressed in the ovary (Flybase 2024). However, its function remains unknown. If ADMA-histones play an important role in the local recruitment of Rhino to the OSC genome, it is highly likely that the loss of DARTs responsible for ADMA modification would influence the subcellular localization of Rhino.

A previous study showed that HP1a germline knockdown (GLKD) causes Rhino foci to be more prominent in the ovary and reduces Rhino occupancy at the clusters^37^. Because a similar phenotype was observed for Rhino foci in HP1a-knockdown (KD) OSCs (Fig. 3A, B), we used this phenotype to examine the loss-of-function effects of DARTs. Similar phenotypes were observed in DART1– and DART4-KD OSCs as in HP1a-KD OSCs, suggesting that DART1/4-dependent histone ADMA modification is involved in Rhino recruitment (Fig. 3A, B). Moreover, the localization of Rhino in DART1/4 KD cells closely matches prominent H3K9me3 foci, suggesting that the arginine methylation activity of DART1/4 contributes to the genomic localization of Rhino in OSCs as a genome scaffold beyond H3K9me3 (Supplementary Fig. 3A). In contrast, this phenotype was not observed for HP1a, suggesting that the effects of DART1/4 are specific to Rhino (Supplementary Fig. 3B).

**Figure 3.**
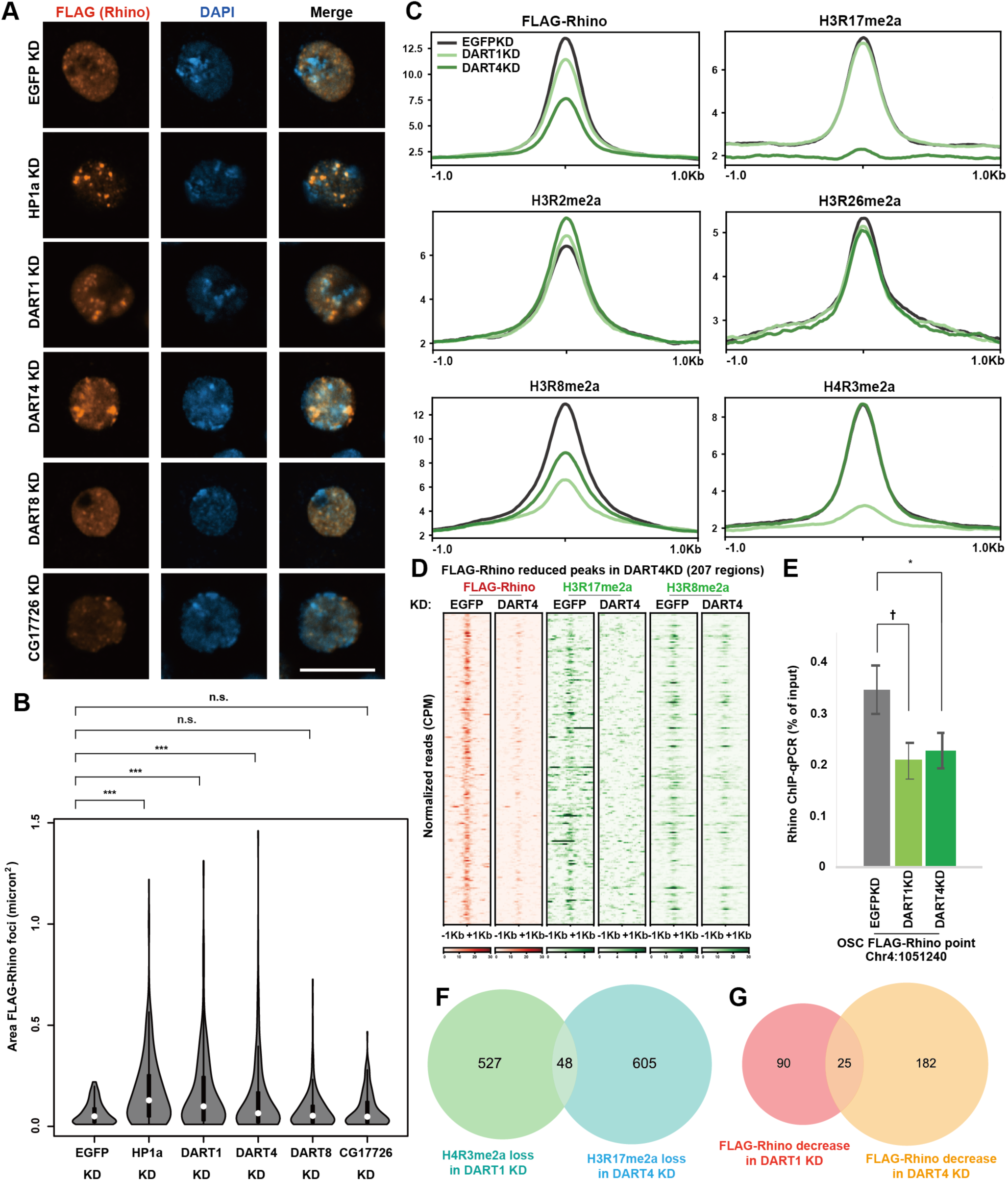
Arginine methyltransferases DART1 and DART4 play a role in Rhino genomic localization and foci formation in OSCs. (A) Confocal immunofluorescent images of FLAG-Rhino (red) and nuclei (blue; DAPI stained) in the EGFP-, HP1a-, DART1-, DART4-, DART8-, and CG17726-KD OSCs. Scale bar represents 10 µm. (B) Comparison of FLAG-Rhino foci sizes among the EGFP-, HP1a-, DART1-, DART4-, DART8-, and CG17726-KD OSCs in (A). Thirty OSCs from three biological replicates were analyzed. Statistical significance in differences of Rhino foci between the genotypes was examined using Kolmogorov-Smirnov test. *P < 0.05, **P < 0.01, and ***P < 0.001. HP1a KD: ***P = 8.49 × 10^−13^, DART1 KD: ***P = 1.83 × 10^−7^, DART4 KD: ***P = 2.83 × 10^−4^, DART8 KD: n.s. P = 0.655, CG17726 KD: n.s. P = 0.454. The box indicates the lower (25%) and upper (75%) quartiles, and the open circle indicates the median. Whiskers extend to the most extreme data points. (C) ChIP-seq profile showing the localization of FLAG-Rhino, H3R2me2a, H3R8me2a, H3R17me2a, H3R26me2a, and H4R3me2a in EGFP-, DART1-, and DART4-KD OSCs at FLAG-Rhino localization sites. The genomic regions used to draw the profile are the same as in Fig. 1G. (D) Heatmap showing the localization of FLAG-Rhino, H3R17me2a, and H3R8me2a in EGFP– and DART4-KD OSCs at the sites of decreased FLAG-Rhino localization in DART4 KD. The regions with decreased FLAG-Rhino were defined as those with an M value greater than 0.5 by manorm. macs2 callpeak p = 0.05 was used for manorm with input as a control. (E) FLAG-Rhino ChIP-qPCR results of one site at which Rhino initially localized. DART1 KD: †P = 5.88 × 10^−2^, DART4 KD: * P = 3.17 × 10^−2^. (F) Venn diagram showing the overlap of H3R17me2a localization decreased in DART4 KD and H4R3me2a localization decreased in DART1 KD. The regions with decreased ADMA-histones were defined as those with an M value greater than 1 in manorm. macs2 callpeak p = 0.01 was used for manorm. (G) Venn diagram showing the overlap of Rhino localizations that were decreased by DART1 KD and DART4 KD. The decreased genomic sites are identified as in (F).

Fluctuation in the localization of Rhino in the DART1– and DART4-KD OSCs was also observed in ChIP experiments. In ChIP-seq, Rhino tended to be decreased in DART1 KD and DART4 KD (Fig. 3C, D), which was confirmed by ChIP-qPCR (Fig. 3E). Among the ADMA-histones that co-localized with Rhino, H3R17me2a decreased in a DART4-dependent manner and H4R3me2a decreased in a DART1-dependent manner (Fig. 3C), consistent with previous findings for human PRMT1 and CARM1^38,39,40,41^. Detailed analysis revealed that the genomic localizations of H4R3me2a lost in DART1 KD (575 sites) and H3R17me2a lost in DART4 KD (653 sites) overlapped at 48 sites (Fig. 3F). Correspondingly, the genomic localizations of Rhino that decreased in DART1 KD (115 sites) and those that decreased in DART4 KD (207 sites) overlapped at 25 sites (Fig. 3G). These results suggest that DART1 and DART4 could contribute to Rhino recruitment at distinct genomic sites through the decreases in ADMA-histones in each of their KD conditions (H4R3me2a and H3R17me2a, respectively).

Among ADMA-histones, only H3R8me2a tended to decrease in both DART1 and DART4 KD (Fig. 3C). To examine the specificity of the regulation of H3R8me2a by DART1 and DART4, the degree of overlap was visualized in Venn diagrams for H3R8me2a, as well as H3R17me2a and H4R3me2a. The genomic localizations of H3R8me2a decreased by DART1 KD (235 sites) and DART4 KD (58 sites) weakly overlapped (8 sites) (Supplementary Fig. 3C). The genomic localizations of H3R8me2a (58 sites) and H3R17me2a (652 sites) that decreased in DART4 KD overlapped only at 7 sites (Supplementary Fig. 3D), as was the case for H3R8me2a (235 sites) and H4R3me2a (575 sites), which were decreased in DART1 KD (Supplementary Fig. 3E). These results suggest that DART1 and DART4 regulate Rhino localization through H3R8me2a at different sites, as well as through H3R17me2a and H4R3me2a.

The level of Rhino expression was unchanged in DART1/4-KD cells (Supplementary Fig. 3F). Rhino and HP1a even in normal OSCs were under detection levels with anti-ADMA antibodies (Supplementary Fig. 3G). Thus, the loss of DART1/4 does not appear to affect Rhino or HP1a function directly through methylation. Although DART4 has been reported to be highly expressed in the ovary^28^, public RNA-seq data showed that its reads were almost undetectable in both OSCs and ovary (Supplementary Fig. 3H). Consistent with this, qPCR analysis revealed that DART4 mRNA expression was only about 3% of that of DART1 (Supplementary Fig. 3I). Nevertheless, this low-level mRNA expression appears sufficient to support its functional role, such as the modification of H3R17me2a (Fig. 3C and Supplementary Fig. 3F). The knockdown of each gene in OSCs was confirmed to be effective (Supplementary Fig. 3J).

### ADMA-histones co-localize with Rhino in a manner dependent of DART4 in the ovary

We examined the localization of ADMA-histones in the Oregon-R ovary. In the wild-type (WT) ovary, we detected Rhino peaks in FLAG-Rhino localization sites in OSCs and ADMA-histones, H3R8me2a, H3R17me2a, and H4R3me2a, co-localized with them (Fig. 4A and Supplementary Fig. 4). Heatmap showed that these ADMA-histones are detected in the narrow Rhino-positive regions as in the case of Rhino in OSCs (Fig. 4B). As well, the metaplot showed that ADMA-histones tend to localize to the ends of piRNA clusters (Fig. 4C). The trend of Rhino genome binding observed in OSCs was also recapitulated in ovaries at corresponding genomic locations.

**Figure 4.**
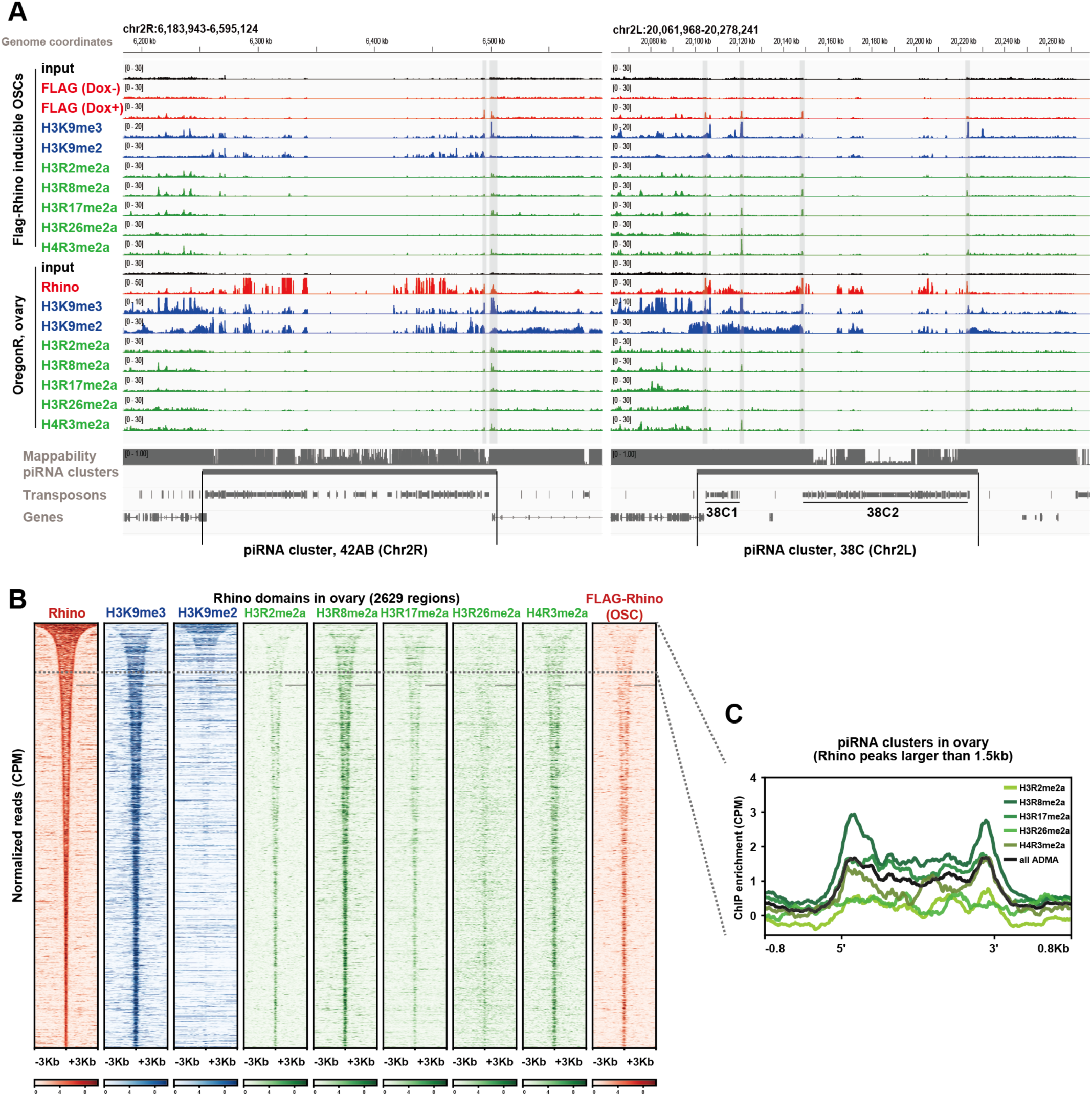
ADMA-histones co-localize with Rhino prior to Rhino spread in the ovary. (A) Genome browser snapshot showing the localization of H3R2me2a, H3R8me2a, H3R17me2a, H3R26me2a, and H4R3me2a in piRNA clusters in OSCs and Oregon-R ovary. The gray-shaded areas indicate the localization sites of FLAG-Rhino in each piRNA cluster in OSCs. (B) Heatmap showing the localization of H3R2me2a, H3R8me2a, H3R17me2a, H3R26me2a, and H4R3me2a at Rhino peaks in Oregon-R ovary. The Rhino peak regions shown in Fig. 1C were used. (C) Metaplot showing the localization of ADMA-histones in the ovary at piRNA clusters. For piRNA clusters, broad peaks of Rhino localization in the ovary greater than 1.5 kb were used.

To determine whether the destabilization of Rhino genome binding observed in OSCs under DART4-KD conditions is also observed in the ovary, we performed GLKD of DART4 using the GAL4-UAS system. Under the DART4-GLKD conditions (Supplementary Fig. 5A), Rhino foci were more prominent in the absence of DART4 (Fig. 5A, B). ChIP-seq experiments showed that Rhino localizes almost the same genomic regions in the ovary as in OSCs, although these localizations are weaker compared to piRNA clusters such as 42AB (Fig. 5C, D). Notably, Rhino levels significantly decreased with the loss of H3R17me2a under DART4-GLKD conditions (Fig. 5C-F, and Supplementary Fig. 5C, D). These results indicate that the relationship between ADMA-histones and Rhino found in OSCs is conserved in the ovary.

**Figure 5.**
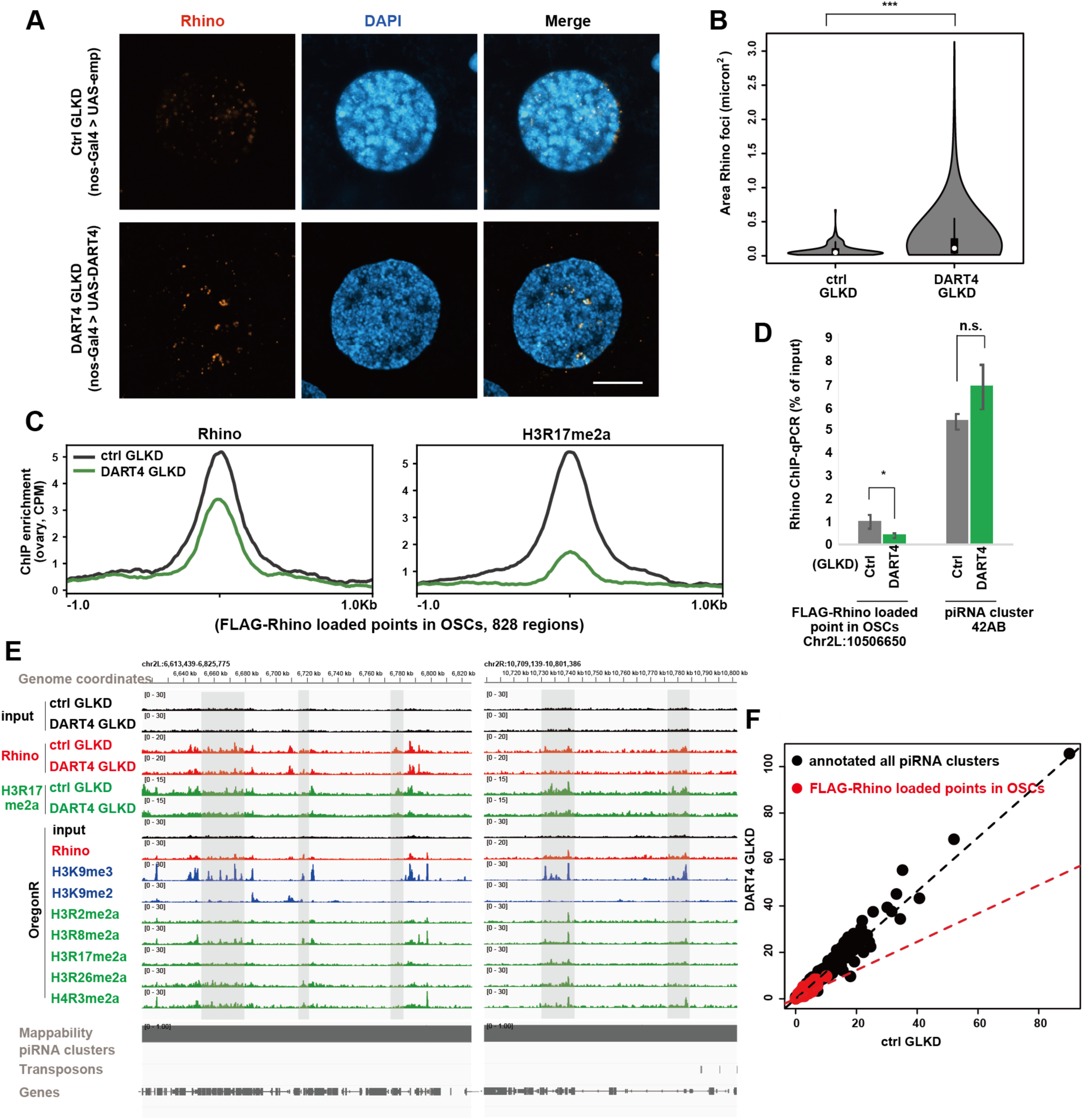
DART4 GLKD has impacts on Rhino foci and Rhino genomic localization in the ovary. (A) Confocal immunostaining images of Rhino (red) in ctrl– and DART4-GLKD ovaries. DAPI shows nuclei (blue). Scale bar represents 10 µm. (B) Comparison of sizes of Rhino foci between the ctrl– and DART4-GLKD ovaries in (A). Ten ovaries from two biological replicates were analyzed. Statistical significance in differences of Rhino foci between the two genotypes was examined as in Fig. 3B. DART4 GLKD: ***P = 1.26 × 10^−10^. (C) ChIP-seq profile showing the localization of Rhino and H3R17me2a in ctrl– and DART4-GLKD ovaries at the FLAG-Rhino localization sites in OSCs. The FLAG-Rhino peak regions, as in Fig. 1G, are used. (D) Rhino ChIP-qPCR results of one site at which Rhino initially localized and piRNA cluster 42AB. t-test was used for statistical hypothesis testing. Rhino initial localization point: * P = 4.03 × 10^−2^, piRNA cluster 42AB: n.s. P = 0.11. (E) Genome snapshot of ChIP-seq data in euchromatic regions. The gray-shaded areas indicate genomic locations where Rhino localization is decreased in a H3R17me2a-dependent manner. (F) Scatter plot comparing Rhino enrichment in ChIP-seq between ctrl GLKD and DART4 GLKD. Each dot represents the square root of the read count per million mapped reads (RPM) within the annotated piRNA clusters in piRNAcluster database^42^ (colored in black, 184 clusters, without clusters on chromosome Y and uni-strand piRNA clusters flamenco/20A) and FLAG-Rhino loaded points in OSCs as in Fig. 5C (colored in red, 828 regions). The dotted lines represent the least-squares linear regression fit to the data points: the black line for 184 piRNA clusters and the red line for 828 FLAG-Rhino–loaded points in OSCs.

Interestingly, Rhino spreading across clusters annotated in the piRNA cluster database^42^, such as 42AB, 38C, 80F, and 102F, was little affected by DART4 GLKD (Supplementary Fig. 5D). These results suggest that the weak Rhino localization dependent on ADMA-histones (such as at the terminal sites of piRNA clusters) and the strong Rhino localization within the core regions of piRNA clusters are fundamentally governed by distinct regulatory mechanisms.

Loss of Kdm3 in the ovary (i.e., Kdm3-GLKD ovary) causes an increase in H3K9me2 and regional expansion, accompanied by Rhino spreading and the appearance of ectopic piRNA clusters^13^. To determine whether ADMA-histones are involved in this process, we examined the localization of ADMA-histones in the ectopic piRNA clusters. ADMA-histones were found near the 5′ and 3′ ends of the Kdm3 GLKD-dependent piRNA clusters, as well as in the authentic piRNA clusters (Supplementary Fig. 5E). Relatively few ADMA-histones were observed inside the clusters, even in the Kdm3 GLKD-dependent piRNA clusters (Supplementary Fig. 5F). This raises the possibility that ADMA-histones may be involved in the Rhino propagation by defining the spreading regions.

### Genomic regions where Rhino binds broadly in a DART4-dependent manner, but not stably anchored, produce some piRNAs

Counting the genomic sites where H3R17me2a was no longer localized in DART4-GLKD ovaries, 3,082 sites were identified. The genomic sites where Rhino was decreased under the same circumstance were also counted (2,078 sites). Of these, 497 sites overlapped in the two groups (Fig. 6A). This indicates that 76.1% of the Rhino genomic localization reduced by DART4 GLKD appears to be H3R17me2a-independent. Forty-three such regions were found in the *Drosophila* genome (Fig. 6B, C). Representative examples are shown in Fig. 6D and Supplementary Fig. 6A: In control KD, these regions have Rhino in their internal regions and H3R17me2a at their ends (and other ADMA-histones at each end in WT Oregon-R strain, albeit at low levels). In DART4 GLKD, H3R17me2a disappeared from the terminal regions as expected. The internal Rhino also disappeared, although the areas did not contain H3R17me2a (Fig. 6D, E). These regions are distinctly different in nature from authentic piRNA clusters, where Rhino localization remained largely unaffected by DART4 GLKD (Fig. 6B).

**Figure 6.**
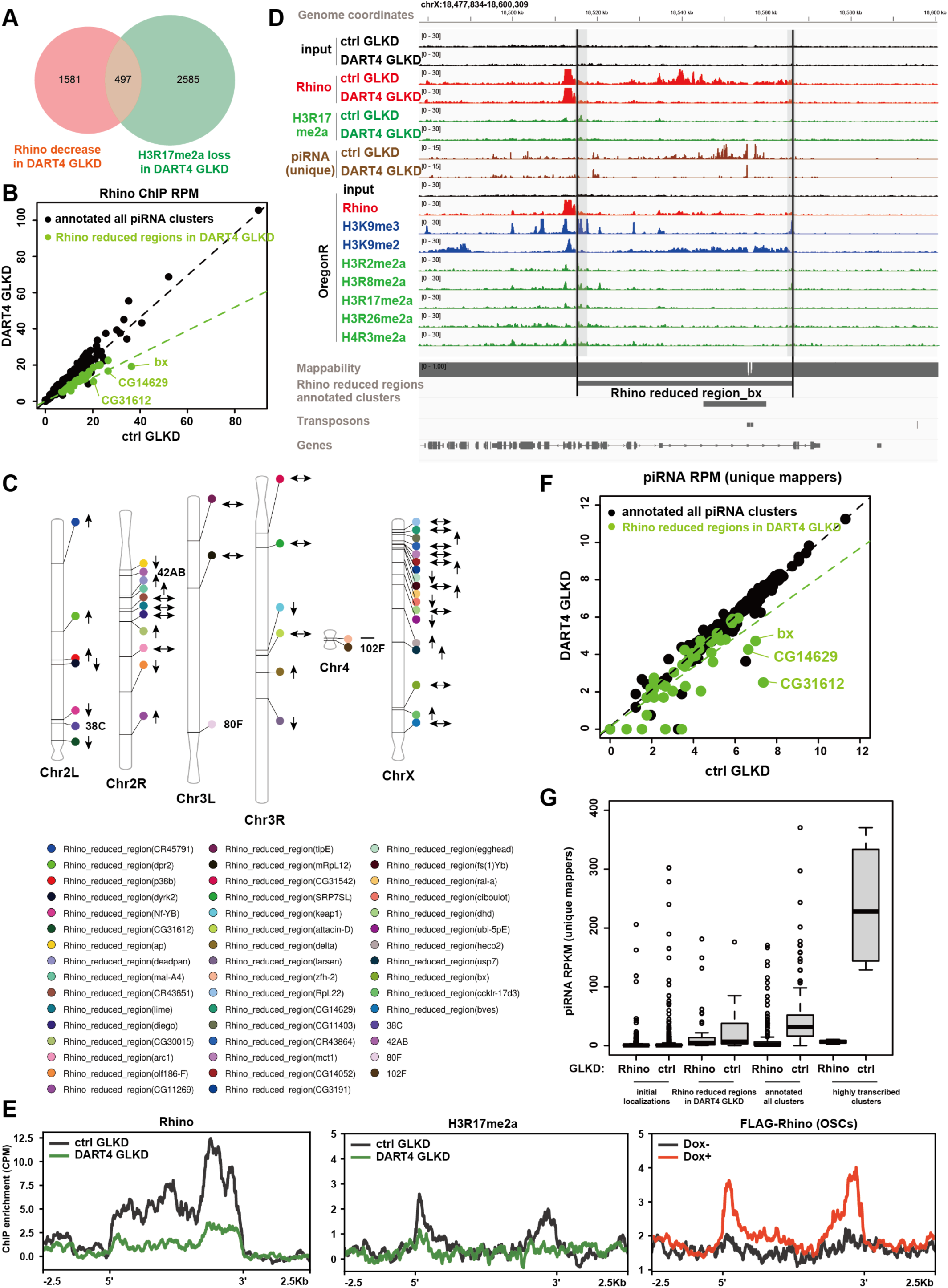
DART4-dependent piRNA source loci (DART4 piSL) are genomic loci where Rhino spreads from H3R17me2a. (A) Venn diagram showing the overlap between sites with reduced Rhino localization and sites where H3R17me2a was lost in DART4 GLKD. The regions where Rhino was decreased were defined as those with an M value greater than 0.5 by manorm, and the regions where H3R17me2a was lost were defined as those with an M value greater than 0. For manorm, the output results of macs2 callpeak p = 0.01 were used with input as a control. (B) Scatter plot comparing Rhino enrichment in ChIP-seq between ctrl GLKD and DART4 GLKD. Each dot represents the square root of the read count per million mapped reads (RPM) within the annotated piRNA clusters in piRNAcluster database^42^ (colored in black, 184 clusters, without clusters on chromosome Y and uni-strand piRNA clusters flamenco/20A) and Rhino reduced regions in DART4 GLKD (colored in green, 43 clusters). The dotted lines represent an approximation by a linear regression model. (C) Chromosomal locations of Rhino reduced regions in DART4 GLKD and the estimated direction of Rhino spread based on initial localization in OSCs. Depressed areas indicate pericentromeres. Arrows show the spread direction based on ADMA-histones. Bidirectional arrows indicate that ADMA-histones are present at both ends and the direction of spread cannot be determined. Horizontal bars indicate clusters without ADMA-histones at both ends. (D) Genome snapshot showing Rhino reduced regions in DART4 GLKD around the *bx* gene. Black lines mark Rhino reduced regions boundaries, and gray-shaded areas highlight potential ADMA-histone localization at cluster ends. (E) ChIP-seq profiles of Rhino and H3R17me2a in ctrl GLKD and DART4 GLKD across all 43 Rhino reduced regions in DART4 GLKD, and Rhino in OSCs. (F) Scatter plot comparing piRNA enrichment between ctrl GLKD and DART4 GLKD. Each dot represents the logged RPM value within piRNA clusters (black: 184 annotated clusters; green: 43 Rhino reduced regions in DART4 GLKD). Dotted lines indicate linear regression. (G) Box plot showing RPKM of piRNAs uniquely mapped to Rhino sites in OSCs, Rhino reduced regions in DART4 GLKD, all piRNAcluster database^42^-annotated clusters, and highly transcribed clusters (42AB, 38C, 80F, 102F). Clusters with RPKM below 400 are shown.

Not only H3R17me2a but also other ADMA-histones were found at the ends of these regions, and H3K9me2 accumulated in the internal areas (Supplementary Fig. 6B), consistent with previous studies suggesting H3K9me2 involvement in the Rhino spreading^13^. FLAG-Rhino in OSCs was also localized at the ends of the regions (Fig. 6E). These results raise the possibility that a subset of Rhino localized to genomic regions correlating with ADMA-histones may serve as origins of spreading.

We next investigated the potential for piRNA production from the genomic regions where Rhino propagates in a DART4-dependent manner but does not stably anchor. Under control conditions, piRNAs were detected mapping to these regions, albeit at low levels (Fig. 6D, F, and Supplementary Fig. 6A). Under DART4-GLKD conditions, piRNA levels were significantly reduced, suggesting that piRNA production from these regions is dependent on DART4. Although piRNA production was also influenced by Rhino, the reduction observed upon Rhino loss was much less pronounced compared to annotated piRNA clusters (Fig. 6G).

The expression levels of transposons and genes presumed to be targeted by piRNAs originating from genomic regions showed minimal changes under DART4-GLKD conditions (Supplementary Fig. 6C). Similarly, egg hatching rates remained largely unaffected (Supplementary Fig. 6D). These observations could be attributed to the relatively low levels of piRNAs arising from these regions compared to authentic piRNA clusters (Supplementary Fig. 6E, F), as well as their shorter genomic lengths (Supplementary Fig. 6G). This can also be explained by the fact that piRNA production from authentic clusters remained largely unaffected. Notably, even the functional loss of authentic piRNA clusters in germ cell piRNA production does not usually result in infertility^43^. Based on these findings, which highlight that these genomic regions differ from authentic piRNA clusters in many aspects yet possess the ability to produce piRNAs, we designate them as DART4-dependent piRNA source loci (DART4 piSL).

Kipferl stabilizes the Rhino-genome binding^12^. ChIP-seq using anti-Kipferl antibodies showed that Kipferl localizes to DART4 piSL and decreases in DART4 GLKD like Rhino (Supplementary Fig. 7A, B). These results suggest that, although Kipferl is present at DART4 piSL, the apparent stabilization of Rhino may depend more on DART4 than on Kipferl, unlike at other authentic piRNA clusters.

In Kipferl mutants, Rhino localization is biased toward satellite repeats^12^. In DART4 GLKD, there appears to be no significant difference in Rhino localization (Supplementary Fig. 7C), nor does Rhino become enriched at satellite repeats (Supplementary Fig. 7D).

A recent comprehensive annotation of active piRNA clusters across eight highly inbred *Drosophila* strains classified the clusters into eight levels of evolutionary novelty^44^. We subsequently analyzed the enrichment of ADMA-histones, which significantly correlate with Rhino localization in OSCs (Fig. 2A-E) and ovaries (Fig. 4 A-C) (e.g., H3R2me2a, H3R8me2a, H3R17me2a, and H4R3me2a), around these piRNA clusters. Notably, a trend of greater ADMA-histone enrichment was observed at the ends of the newer clusters (Fig. 7A). This trend was also observed in DART4 piSL (Supplementary Fig. 8A). In contrast, such bias was not observed in older clusters or other genomic loci (Fig. 7A, B, Supplementary Fig. 8A). H3R2me2a, H3R8me2a, H3R17me2a, and H4R3me2a showed a loss of enrichment as the clusters aged (Fig. 7A). The DART4-dependent H3R17me2a differences observed between newer and older clusters were less pronounced in DART4 GLKD (Fig. 7B). We next extracted piRNA clusters where piRNAs were detected in the Oregon-R strain and examined them for ADMA-histone enrichment. The results remained consistent (Supplementary Fig. 8B). A similar analysis using GLKD strains yielded identical findings (Supplementary Fig. 8C). These results suggest that DART4 piSL represent relatively new piRNA clusters.

**Figure 7.**
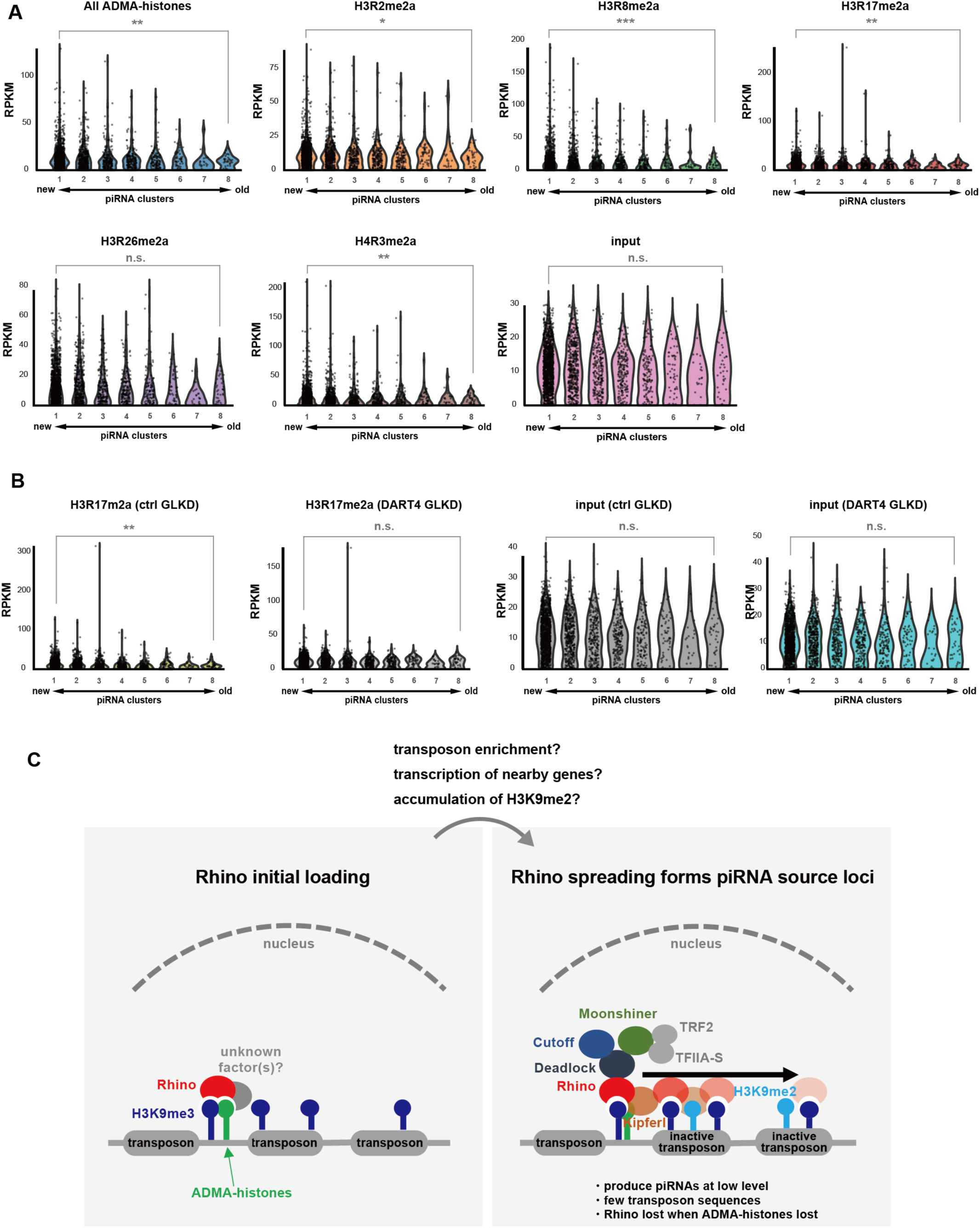
ADMA-histones are enriched at the ends of evolutionarily young piRNA clusters. (A) RPKM of each ChIP-seq dataset (Oregon-R) in the piRNA cluster dataset with eight levels of evolutionary novelty. RPKM was calculated for the 500 bp region surrounding the ends of the piRNA clusters. t-test was used for statistical hypothesis testing between clusters found only in 1 strain (cluster 1, newly emerged) and clusters found in all 8 strains (cluster 8, evolutionarily conserved). All ADMA-histones: ** P = 1.11 × 10^−3^, H3R2me2a * P = 1.45 × 10^−2^, H3R8me2a: *** P = 1.27 × 10^−4^, H3R17me2a: ** P = 7.28 × 10^−3^, H3R26me2a: n.s. P = 0.23, H4R3me2a: ** P = 1.91 × 10^−3^, input: * P = 0.53. (B) Same analysis of Fig. 7A for each ChIP-seq dataset (ctrl GLKD and DART4 GLKD). H3R17me2a (ctrl GLKD): ** P = 2.77 × 10^−3^, H3R17me2a (DART4 GLKD): n.s. P = 0.24, input (ctrl GLKD): n.s. P = 0.42, input (DART4 GLKD): n.s. P = 0.47. (C) Model for the piRNA source loci formation. In the cell conditions lacking Kipferl such as in OSCs, Rhino initially localizes to bivalent nucleosomes containing H3K9me3 and ADMA-histones (left). In the second step, Rhino spreading occurs, leading to the formation of DART4 piSL. Transcription of adjacent regions and/or H3K9me2 enrichment may play a role in this process (right). Rhino in DART4 piSL is lost if ADMA-histones are lost. It remains unclear whether DART4 piSL can eventually develop into highly transcribed piRNA clusters, such as 42AB, over the course of evolutionary time.

Based on all these results, we propose that ADMA-histones play a crucial role in determining Rhino’s initial genomic binding (Fig. 7C, left). Furthermore, ADMA-histone-dependent Rhino binding to the genome may also contribute to the propagation of Rhino within the piRNA cluster regions (Fig. 7C, right). Our evolutionary analysis suggests that DART4 piSL appear to share features with young piRNA clusters and may eventually develop into *bona fide* piRNA clusters like 42AB; however, it does not provide direct evidence for this process. Further studies will be necessary to explore this possibility.

## Discussion

In this study, we took advantage of cultured OSCs for our analysis and found that chromatin marks (i.e., bivalent nucleosomes containing H3K9me3 and ADMA-histones) appear to contribute to the initial loading of Rhino onto the genome. Comparative analysis of OSC and ovarian ChIP-seq data also allowed us to identify “primitive” piRNA source loci, DART4 piSL. Based on these results, DART4 piSL are piRNA source loci where Rhino propagates in a DART4-dependent manner from the ADMA marks at the ends, but this spread is not stable. In DART4 piSL, Rhino spreads from the initial loading sites but does not stabilize, indicating that these regions may be nascent piRNA clusters. The evolutionary analysis showing a stronger tendency for ADMA accumulation at the ends of younger piRNA clusters (Fig. 7A) provides indirect evidence that DART4 piSL may represent evolutionarily young, nascent piRNA clusters. While DART4 piSL may represent a “later” stage compared to annotated piRNA clusters, the essential role of piRNAs in reproduction and species conservation makes it unlikely that a large number of clusters exist in a degenerate state. Thus, we speculate that these piRNA source loci represent an “earlier” stage in piRNA cluster formation. As note, both ctrl GLKD and DART4 GLKD share the nos-GAL4 background, but we cannot fully exclude the possibility that the identified DART4 piSL reflect strain-specific genomic variation between ctrl UAS-RNAi strain and DART4 UAS-RNAi strain rather than DART4-dependent regulation.

Whether DART4 piSL evolve into large piRNA clusters such as 42AB and 38C during the course of evolution remains an open question. Although these large clusters were not affected by DART4 GLKD, ADMA-histones were still detectable, albeit at low levels, at the ends of their Rhino peaks (Fig. 2B and 4A). This observation raises the possibility that even large piRNA clusters like 42AB and 38C may have initially formed through a process involving ADMA-histone–mediated Rhino loading followed by propagation, as seen in DART4 piSL. While it remains possible that these clusters formed through a different mechanism without passing through a DART4 piSL-like phase, such a scenario would leave the presence of residual ADMA-histones at their ends unexplained. Nonetheless, evolutionary analyses provide only indirect evidence, and experimental validation will be necessary to determine whether DART4 piSL can indeed mature into larger piRNA clusters.

In general, ADMA-histones are known to function as transcriptional coactivators^45^. In mammalian cells, H3R8me2a does not co-localize with H3K9me3, but with enhancer modifications such as H3K4me1 and H3K27ac^25^. This may indicate that the function of ADMA-histones has evolved in a *Drosophila*-specific manner, which is not unlikely given that Rhino itself is *Drosophila*-specific. However, there are regions where Rhino does not localize in the read sets of ADMA-histones and H3K9me3, suggesting that an unknown factor may contribute to the specificity of Rhino targeting (Fig. 2E).

This study showed that multiple ADMA-histones, such as H3R17me2a and H4R3me2a, play a role in the initial Rhino loading. This suggests that a co-factor(s) with an affinity for ADMAs might be involved. Tudor domains are known to bind partner proteins through methylation of arginine and lysine residues^46^, and Tudor domain-containing protein 3 (TDRD3) was identified as a factor that specifically binds to H3R17me2a and H4R3me2a in humans^47^. Therefore, TDRD protein family members may be involved in Rhino loading via ADMAs. It should also be noted that not all ADMA-enriched regions recruit Rhino; rather, Rhino association is observed only where ADMA-histones overlap with H3K9me3 (Fig. 2E). Even the combination of H3K9me3 and ADMA-histones does not fully explain the specificity of Rhino recruitment, implying the requirement for additional co-factors. Such factors may include other ADMA marks (e.g., H3R42me2a) or chromatin-interacting proteins that are arginine-methylated by DART1 or DART4. DART1 and DART4 might contribute to Rhino recruitment not only through direct histone methylation but also via arginine methylation of non-histone chromatin factors. However, at least under our conditions, HP1a methylation did not appear to be involved in Rhino behavior (Fig. S3G). Given the potential functional redundancy among multiple DARTs, future studies may require simultaneous knockdown or knockout of several DART proteins to fully assess their contribution to Rhino recruitment and piRNA cluster regulation.

The mechanism by which ADMA-histones is involved in Rhino propagation remains unclear. Because Rhino propagation has been shown to be correlated with transcription^7,10^, one interpretation is that ADMA-histones may affect Rhino spreading via transcriptional activity. Insertion of insulators into piRNA clusters inhibits piRNA production^48^, indicating that the three-dimensional (3D) structure of the genome within the piRNA cluster also affects the extent of Rhino spreading. ADMA-histones were found at the ends of piRNA clusters, reminiscent of the boundaries of topologically associated domains. An unknown stabilizing factor(s) suggested by the absence of Kipferl in some DART4 piSL (Supplementary Fig. 7A, B) may be a relevant factor(s) determining such genomic 3D structures. Functional validation of ADMA-histones in piRNA cluster specification will be required in future studies.

Maternally inherited siRNAs initiate piRNA biogenesis from a transgene inserted into the uni-strand piRNA cluster 20A^14^. In this previous report, evidence was presented that siRNAs induce “*de novo”* Rhino loading onto this transgene (20A-X) via H3K9me3. This led us to hypothesize that ADMA-histones should be present in the vicinity of 20A-X. Indeed, we detected ADMA-histones in the region and Rhino, albeit weakly (Supplementary Fig. 4). Thus, this study provided experimental evidence in support of the “*de novo* model” by reinforcing the findings of siRNA induced piRNA biogenesis. This model has been becoming supported by evolutionary analyses^44,48,49^, but experimental results had been limited. The evolutionarily young piRNA clusters, which appear in a limited number of strains, may have formed through the spread of Rhino into newly inserted transposons, facilitated by the presence of ADMA-histones in their vicinity (Fig. 7A).

In *Drosophila*, major piRNA clusters such as 42AB and 38C are located at the boundary between heterochromatin and euchromatin^4,51^. It was unclear why these highly transcribed clusters, which are capable of producing substantial piRNAs, tend to appear in regions where two different types of chromatins collide. If DART4 piSL represent the initial stage in the formation of piRNA clusters, our model may help explain this question by showing that cluster formation begins with initial Rhino loading by ADMA-histones onto euchromatic regions and then spreads to adjacent regions of H3K9me2-rich heterochromatin (Fig. 7C). Rhino shows a tendency to spread from an DART4 piSL on chromosome 2 to heterochromatin/euchromatin boundary regions where 42AB and 38C are present (Fig. 6C), supporting our hypothesis.

## Methods

### Establishment and cloning of FLAG-Rhino inducible OSCs

To construct Tet-ON 3G inducible FLAG-Rhino plasmid, cDNA encoding FLAG-Rhino was cloned into pPB-2× Ty1-Tjen-EGFP-P2A-BlastR^52^ using In-Fusion cloning (TAKARA), bearing pPB-FLAG-Rhino-Tjen-EGFP-P2A-BlastR. The Actin promoter and *EGFP* cDNA were replaced respectively with the TRE3G promoter and *rtTA* cDNA from inducible Caspex expression plasmid (Addgene). PCR was performed using PrimeSTAR MAX (TAKARA) with primers (Supplementary Table 1). FLAG-Rhino inducible OSCs were established using the piggyBac system as described previously^52^. In brief, 2 × 10^6^ OSCs were transfected with 4 μg of pPB-TRE3G-FLAG-Rhino-Tjen-trTA-P2A-BlastR and 1 μg of pHsp70-Myc-hyPBase using Nucleofector Kit V (Lonza). After incubation in media containing blasticidin (50 μg/mL), colonies were isolated. One of the colonies contained a single FLAG-Rhino copy in the intron of the *jing interacting gene regulatory 1* gene, and this was used in this study.

### Cell culture and RNAi

FLAG-Rhino inducible OSCs were cultured in Shields and Sang M3 Insect Medium (Sigma) supplemented with 10% fly extract, 10% fetal bovine serum, 0.6 mg/mL glutathione, and 10 mU/mL insulin^26,53^. To perform RNAi, 400 pmol of siRNA duplex was introduced into 3.0 × 10^6^ cells using the Cell Line 96-well Nucleofector Kit SF (Lonza) with program DG150 of the 96-well Shuttle Device (Lonza). After 48 h incubation, cells were transfected with another 400 pmol of siRNA and incubated further for 48 h. For HP1a KD, RNAi was performed once to avoid cell damage. All siRNAs used are shown in Supplementary Table 2.

### Fly stock

Flies were raised in standard Bloomington medium at 26 °C. Oregon-R was employed as a WT strain. For DART4 GLKD, BDSC 4937 (nosGAL4, w[1118]; P{w[+mC]=GAL4::VP16-nanos.UTR}CG6325[MVD1]) and BDSC 36833 (y[1] sc[*] v[1] sev[21]; P{y[+t7.7] v[+t1.8]=TRiP.GL01073}attP2) were used as the maternal GAL4 driver and DART4 UAS-RNAi lines, respectively. BDSC 36303 (y[1] v[1]; P{y[+t7.7]=CaryP}attP2) was used as a control (ctrl).

### Western blotting

Western blotting was performed as described previously^54^. Antibodies are summarized in Supplementary Table 3. Nuclear extraction and immunoprecipitation were performed as described previously^55^.

### Immunofluorescence

Immunofluorescence of OSCs was performed essentially as described previously^56^. OSCs were fixed using 4% formaldehyde (Wako) and permeabilized in PBS containing 0.1% Triton X-100 for 24 h at RT. For DAPI staining, VECTASHIELD Mounting Medium with DAPI (Vector Laboratories) was used. Immunofluorescence of ovaries was performed as described previously^10^. All images were obtained using a confocal laser scanning microscope (Olympus FV3000 or Carl Zeiss LSM 710). For image processing, ZEN software (Carl Zeiss) was used. Antibodies used are listed in Supplementary Table 3.

### Generation of mouse anti-Rhino monoclonal antibody

Anti-Rhino monoclonal antibody for immunofluorescence in the ovary were raised against N-terminal 200 residues of Rhino using a protocol published previously^57^. The N-terminal 200 residues of Rhino were fused to glutathione S-transferase (GST) and the purified fusion protein was used as antigen to immunize mice. The expression vector was produced by inserting the cDNA encoding the Rhino fragment into pGEX-5X-3 (Cytiva) by Infusion (Takara). The primers used are listed in Supplementary Table 1. The specificity of the antibody was validated using OSCs (Supplementary Fig. 9A).

### RNA extraction and RT-qPCR

Total RNAs were extracted from OSCs or ovaries with the ISOGEN II reagent (Nippon Gene) and treated with DNase (Life Technologies) to eliminate DNA contaminations. The cDNA libraries were prepared using the Transcriptor First Strand cDNA Synthesis Kit (Roche). For RT-qPCR, cDNAs were amplified using the StepOnePlus Real-Time PCR System (Applied Biosystems) with PowerUp SYBR Green Master Mix (Thermo Fisher Scientific). The expression level of target genes and transposons in OSCs and ovary were calculated by ΔΔCt method using *ribosomal protein 32* (*RP49/RpL32*) as an internal control. Data are presented as mean values ± SE. The primers used are listed in Supplementary Table 1.

### OSC ChIP

OSCs were fixed in PBS containing 1% formaldehyde for 10 min at RT and glycine was added to 150 mM to stop the fixation. Cells were collected and suspended in swelling buffer [25 mM HEPES-KOH (pH 7.3), 1.5 mM MgCl_2_, 10 mM KCl, 1 mM DTT, 0.1% NP-40, supplemented with cOmplete ULTRA Tablets, EDTA-free (Roche)]. The nuclei were collected by centrifugation at 1,000 g for 5 min at 4°C, suspended in sonication buffer [50 mM HEPES-KOH (pH 7.4), 140 mM NaCl, 1 mM EDTA, 1% Triton X-100, 0.1% sodium deoxycholate, 0.1% SDS, supplemented with cOmplete ULTRA Tablets, EDTA-free], and sonicated using a Covaris S220 Focused ultrasonicator for 10 min at 4°C with setting of peak power of 200, duty factor of 10.0, and 200 cycles/burst. The chromatin fraction was obtained by centrifugation at 20,000 g for 15 min at 4°C and diluted with sonication buffer to make the DNA concentration to 0.2 mg/mL. Immunoprecipitation was performed O/N at 4°C using 1mL of chromatin fraction per ChIP. Antibodies used are listed in Supplementary Table 3. The reaction mixture was mixed with 25 μL of Dynabeads ProteinG (Invitrogen) and incubated for 1 h at 4°C. Beads were washed one time each with low-salt buffer [20 mM Tris-HCl (pH 8.0), 150 mM NaCl, 2 mM EDTA, 0.1% Triton X-100, 0.1% SDS, and 1 mM phenylmethylsulfonyl fluoride (PMSF)], high-salt buffer [20 mM Tris-HCl (pH 8.0), 500 mM NaCl, 2 mM EDTA, 0.1% Triton X-100, 0.1% SDS, and 1 mM PMSF], and LiCl buffer [20 mM Tris-HCl (pH 8.0), 250 mM LiCl, 2 mM EDTA, 1% NP-40, 1% sodium deoxycholate, and 1 mM PMSF], and then twice with TE buffer [20 mM Tris-HCl (pH 8.0), 1 mM EDTA, and 1 mM PMSF]. The beads were suspended in elution buffer [50 mM Tris-HCl (pH 8.0), 10 mM EDTA, and 1% SDS] and incubated at 65°C for 30 min. For de-crosslinking, the supernatant was incubated O/N at 65°C after supplement of NaCl to 200mM. RNAs and proteins were eliminated from the sample by incubating at 37°C for 30 min with 10 mg/mL RNase A and 20 mg/mL proteinase K at 55°C for 1 h. The DNA in the supernatant was purified by EconoSpin column IIa (Funakoshi). 50 μL of nuclease-free water was added to it and the sample was centrifuged at 13,000 rpm for 1 min to obtain ChIP product. The following experiments were conducted to evaluate the specificity of the antibodies involved in the main claims of this paper in ChIP-seq: H3R8me2a: ChIP-seq using two different anti-H3R8me2a antibodies showed that signal was obtained at the same genomic locations (Supplementary Fig. 9B). H3R17me2a: A reduction in DART4-KD cells was demonstrated in separate experiments using western blotting (Supplementary Fig. 9C). H4R3me2a: A histone peptide array kit (Active motif) confirmed that the antibody recognizes the H4R3me2a peptide with high specificity (Supplementary Fig. 9D). The histone peptide array was performed according to the manufacturer’s protocol.

### Ovary ChIP

150 ovary pairs were crosslinked in PBS containing 1% formaldehyde for 10 min at RT, quenched with 180 mM glycine, and washed twice in PBS. Ovaries were flash frozen and suspended with swelling buffer (see “OSC ChIP”). Ovaries were disrupted using a Dounce homogenizer and incubated at 4°C for 10 min. The nuclei were collected by centrifugation at 1,000 g for 5 min at 4°C, suspended in sonication buffer, and sonicated (see “OSC ChIP”). The process after this is as described above (see “OSC ChIP”). Antibodies used are listed in Supplementary Table 3.

### ChIP-seq library preparation

ChIP-seq libraries were constructed using NEBNext Ultra II FS DNA Library Prep Kit for Illumina (New England BioLabs) using ChIP products from “OSC ChIP” and “Ovary ChIP”. In the adaptor ligation step, the original adaptor reagent was diluted 10-fold. Adapter ligated DNAs were PCR-amplified (10-12 cycles) with Illumina primers. The PCR products of 100-1,000 bp (including adaptor and primer sequences) were purified using AMPure XP beads (Beckman Coulter) and captured on an Illumina flow cell. The libraries were sequenced using the Illumina MiSeq.

### Computational analysis of ChIP-seq

Prior to mapping, low-quality reads were removed and adapters were trimmed using Fastp^58^. The filtered reads were mapped to the genome of *Drosophila melanogaster* (dm6) using Bowtie 2^59^. Only uniquely mapped reads were used in the analysis using samtools view command option-q 42^60^. For genome browser view, we mapped these unique reads and the scores of the *Drosophila* genome mappability calculated using GenMap software with the k-mer size set to 150^61^. We performed ChIP-seq experiment using at least two biological replicates, except for ChIP with anti-FLAG antibody in OSCs under Dox = 0.05 and 0.5ng/mL conditions. The replicates were highly correlated, so we merged them together and used the merged libraries for analyses. Because of the low mapping score, four replicates were prepared for Rhino and three for Kipferl in the GLKD condition. Peakcall was completed with MACS2 with q = 0.01 settings using input ChIP-seq data as control unless otherwise noted in figure legends^62^. For FLAG-Rhino peakcall in OSCs, anti-FLAG ChIP-seq data in Dox = 0 ng/mL condition was used as control. For Rhino peakcall in Oregon-R ovary in Fig. 1C, D and Fig. 4B, and H3K9me3 peakcall in OSCs in Fig. 1F, options ––broad ––broad-cutoff 0.01 were used. For Fig. 1D, called peak of more than 1.5 kb were extracted to select piRNA clusters where Rhino was localized broadly. For heatmaps and profile plots, ChIP scores were calculated by deeptools^63^. Bigwig files of ChIP were produced by the bamCoverage / bamCompare program of deeptools and matrix data were calculated by the computeMatrix program of deeptools using bigwig files and plotted. For bamCompare program, anti-FLAG ChIP data with Dox = 0 (ng/mL) was subtracted as control for FLAG-Rhino in OSCs and input for the other data. Mapping file correlation and hierarchical clustering were computed using multibamSummary programs of deeptools. To draw the Venn diagram, the narrowPeak file obtained with Macs2 callpeak was stored in the findOverlapsOfPeaks function in R’s ChIPpeakanno library and visualized with makeVennDiagram program^64^. The manorm was used for quantitative comparison of called peaks^65^. As ChIP-seq quality metrics, aligned rates are provided in Supplementary Table 4, while library complexity values and the SSP (strand cross-correlation peak)^66^, a signal-to-noise ratio that can be calculated independently of read number or peak shape, are listed in Supplementary Table 5.

### ChIP-qPCR

Real-time qPCR was performed using PowerUp SYBR Green Master Mix and StepOnePlus real-time PCR system. Data are presented as mean values ± SE. The primers used are listed in Supplementary Table 2.

### Definition of DART4-dependent piRNA source loci

DART4-dependent piRNA source loci were defined using manorm results based on the macs2 callpeak with option p = 0.01 and ––broad ––broad cutoff 0.1 output files that met the following seven criteria. First, M value > 0.5 is required for Rhino to be predominantly enriched against DART4 GLKD. Second, the sharp peaks of the mononucleosome units were excluded because of their non-spreading initial Rhino localization. Third, peaks overlapped with DART4-dependent H3R17me2a were excluded because these peaks cannot distinguish spread Rhino and initial Rhino loading. Fourth, each Rhino peak was checked in the genome browser, and if there were consecutive Rhino peaks around it that met the first, second and third criteria, they were grouped together as one peak. Fifth, peaks that overlap with major piRNA clusters such as 42AB, 38C, 80F, and 102F were excluded because they are highly transcribed piRNA clusters. Sixth, peaks on subtelomeres and pericentromeres were excluded because ChIP-seq reads cannot distinguish piRNA clusters or repeats. Finally, peaks whose localization spanned the entire length of a single transposon were excluded because they may reflect genomic differences between ctrl GLKD and DART4 GLKD, such as copy number variation of transposons. The combined peaks that met these seven criteria were defined as single DART4 piSL together with the surrounding contiguous ctrl GLKD-specific Rhino localizations if they were more than 5 kb. The genomic location of DART4 piSL in Fig. 6C was visualized using Phenogram software^67^. The regions of DART4 piSL are listed in Supplementary Table 6. The regions where Rhino localization increased upon DART4 GLKD were identified as regions with an M value of less than-2 and are listed in Supplementary Table 7. Genes or transposons matched by piRNAs produced from DART4 piSL were analyzed using bedtools and are listed in Supplementary Table 8.

### Ovary piRNA extraction

100 ovary pairs were disrupted using a Dounce homogenizer and incubated in lysis buffer [20 mM HEPES-KOH (pH7.3), 150 mM NaCl, 1 mM EDTA, 1 mM DTT, 0.1% NP-40, 40 U/ml RNasin (Promega), 2 μg/ml pepstatin, 2 μg/ml leupeptin, 0.5% aprotinin] at 4°C for 10 min. The homogenized ovaries were centrifuged at 13,500rpm for 15 min at 4°C. The supernatant was then collected and cooled on ice. 10µg of anti-Piwi antibody was immobilized on Dynabeads protein G (Invitrogen) in lysis buffer at 4°C for 5 min. Beads were incubated with ovarian lysates at 4°C for 2 h, and then washed three times with low-salt wash buffer [20 mM HEPES-KOH (pH7.3), 300 mM NaCl, 1 mM DTT, 0.05% NP-40, 2 μg/ml pepstatin, 2 μg/ml leupeptin, 0.5% aprotinin], three times with high-salt wash buffer [20 mM HEPES-KOH (pH7.4), 500 mM NaCl, 1 mM DTT, 0.05% NP-40, 2 μg/ml pepstatin, 2 μg/ml leupeptin, 0.5% aprotinin]. After wash, Piwi-bound RNAs were extracted from the beads with phenol–chloroform (Ambion) and then precipitated with ethanol. The precipitated RNAs were denatured with Gel Loading Buffer II (Ambion) for 5 min at 95°C, then electrophoresed on 12% urea-PAGE gel. 20-30 nt nucleic acids were gel purified as piRNAs. piRNA-seq library preparation followed a previous study^68^.

### Computational analysis of piRNA-seq

Raw reads were adapter trimmed and the random 4 bases of each side were removed using cutadapt^69^. Trimmed reads were mapped to the dm6 genome using bowtie with the option-m 1^70^. For read count in Supplementary Fig. 7D, this option was excluded. Read counts for each piRNA cluster were performed using bedtools^71^. For the analysis of piRNA in Rhino GLKD, GSM1580790 and GSM1580793 datasets^72^ were used as piRNA-seq data in the ovaries of ctrl GLKD and Rhino GLKD, respectively.

### Egg hatching assay

50 males and 50 virgin females were collected and mated in a cage with apple juice agar plate and yeast paste for 2 days at 25°C. After exchanging plates twice in the morning, the eggs on the plates were collected within 3 hours. Two hundred eggs were placed on new apple juice agar plates and incubated at 25°C for 2 days, followed by counting the number of hatched larvae.

### Evolutionary analysis of piRNA clusters

The 500 bp around the ends of a novel dataset comprising piRNA clusters categorized into eight levels of evolutionary novelty^44^ (“master-list” of Supplementary Table S4 in reference 44) was used for analysis. The ADMA-histones and input ChIP-seq reads mapped to the piRNA cluster regions were counted by bedtools and reads per piRNA cluster were calculated.Genomic regions with RPKM values above 30 in the input were excluded as they were considered ChIP-seq background noise. For Supplemntary Fig. 8B and 8C, only piRNA clusters with non-zero piRNA RPKM were used for analysis in the Oregon-R (Supplemntary Fig. 8B) and ctrl GLKD (Supplemntary Fig. 8C) strains, respectively. SRR4473614 was used as the piRNA of Oregon-R. Extraction of piRNA from SRR4473614 followed the methods of the original paper^50,73^.

### Computational analysis of RNA-seq

Prior to mapping, low-quality reads were removed and adapters were trimmed using Fastp^58^. The filtered reads were mapped to the genome of *Drosophila melanogaster* (dm6) using STAR^74^ with default settings. Reads of each gene were calculated using featureCount^75^ with default settings.

## Data availability

ChIP-seq and piRNA-seq data generated in this study have been deposited in the Sequence Read Archive (SRA) under accession number PRJNA1085006 and the Gene Expression Omnibus (GSE) under accession number GSE261403. All other data supporting the findings of this study are available from the corresponding authors upon request.

## Acknowledgments

We are grateful to all members of the Siomi laboratory for discussions and comments on this work, especially Kaoru Sato. We also thank Soichiro Yamanaka, Tetsuji Kakutani, and Soichi Inagaki for critical reading of the manuscript, Takashi Fukaya and Hiroshi Kimura for discussion, Yuica Koga for technical assistance with *Drosophila* genetics, and Akashi Taguchi for support with the Illumina sequencing. We are also grateful to Julius Brennecke and Laszlo Tirian for antibodies and Harvard TRiP and Bloomington stock centers for *Drosophila* flies. This study was supported by MEXT KAKENHI Grant Number JP19H05466 and JSPS Grant-in Aid for Transformative Research Areas [A] 25H01303 (to M.C.S.), JSPS KAKENHI Grant Number 21K20629 and 23K14232 (to R.S.), JSPS KAKENHI Grant Numbers 17K08644 and 20H03439 (to K.M.), and JSPS KAKENHI Grant Numbers 24K18093 (to Y.N.)

## Author contributions

R.S. and R.H. conducted biochemical experiments. R.S. and Y.N. performed bioinformatic analysis. H.I. produced FLAG-Rhino inducible OSCs. K.M. produced anti-Rhino antibody. R.S. and M.C.S. conceived the project and designed the experiments. R.S. and M.C.S. wrote the manuscript with input from all authors.

---

**Supplementary Figure 1.**
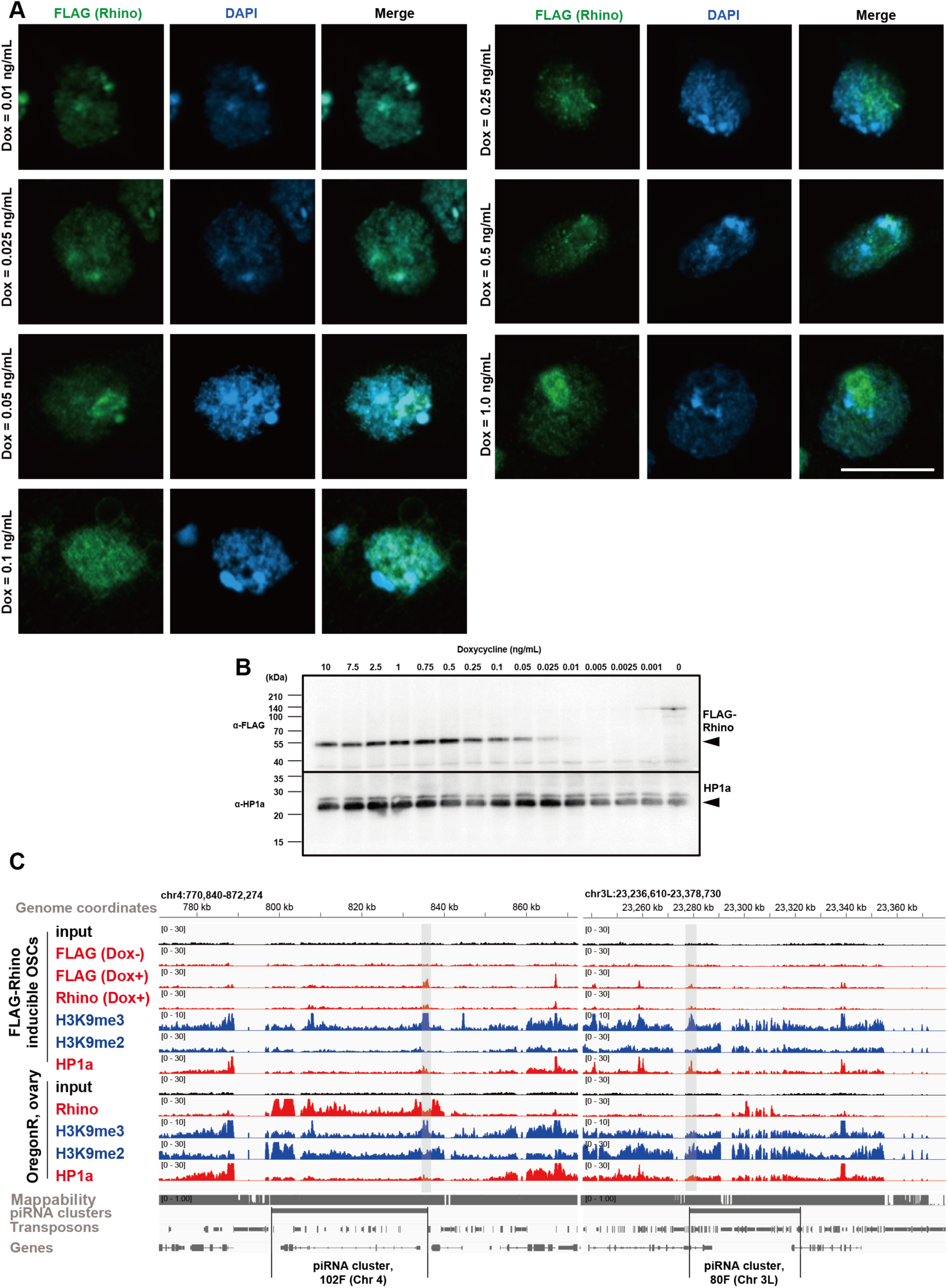
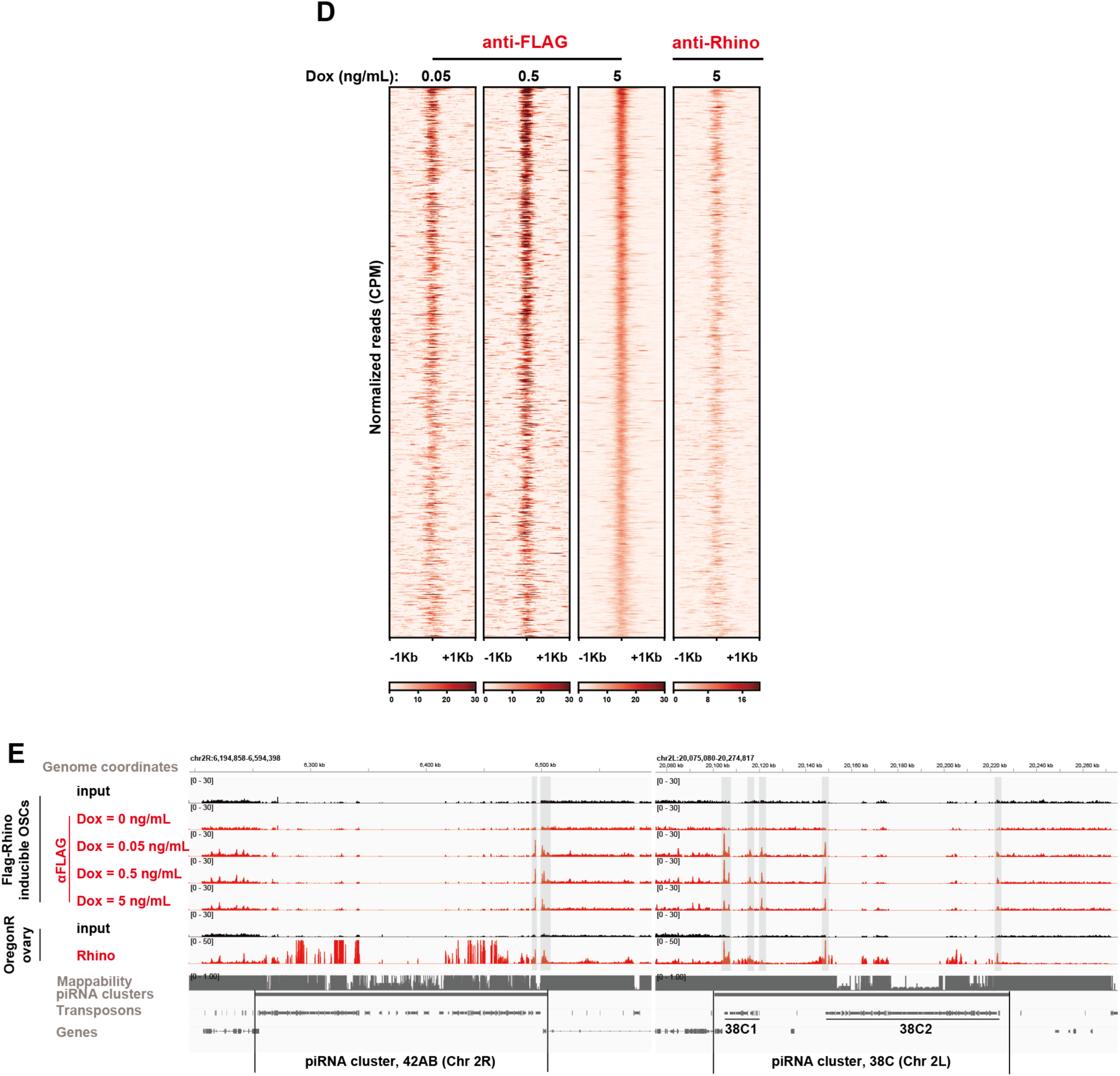
Expression and localization of FLAG-Rhino at different Dox concentrations. (A) Confocal immunofluorescent images of FLAG-Rhino (green) in FLAG-Rhino inducible OSCs upon Dox induction (from 0.01 to 1.0 ng/mL). DAPI shows nuclei (blue). Scale bar represents 10 µm. (B) Western blotting of FLAG-Rhino under each Dox concentration. HP1a was detected as an internal control. (C) Genome snapshot showing ChIP-seq in major dual-strand piRNA clusters (80F and 102F). The gray-shaded areas indicate the sites at which FLAG-Rhino is localized in each piRNA cluster in OSCs. (D) Heatmap showing the localization of FLAG-Rhino in OSCs under various Dox concentrations and Rhino in ovary at FLAG-Rhino peaks of OSCs. The regions of 2 kb around the FLAG-Rhino localization sites were plotted based on macs2 callpeak results. (E) Genome snapshot showing ChIP-seq in major dual-strand piRNA clusters (42AB and 38C). The gray-shaded areas indicate the sites at which FLAG-Rhino is localized in each piRNA cluster in OSCs. piRNA cluster 38C has two piRNA-producing regions, labeled 38C1 and 38C2.

**Supplementary Figure 2.**
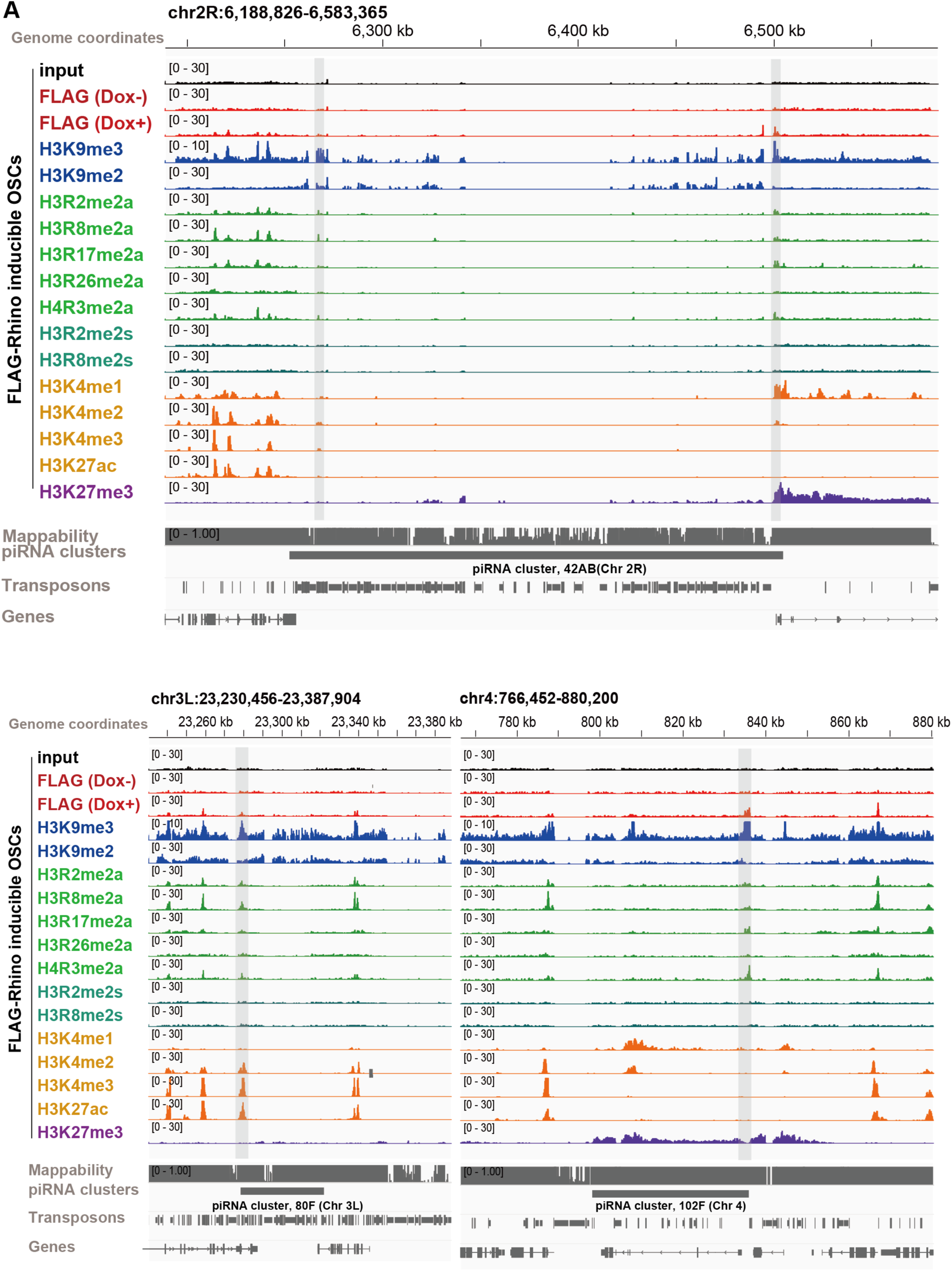
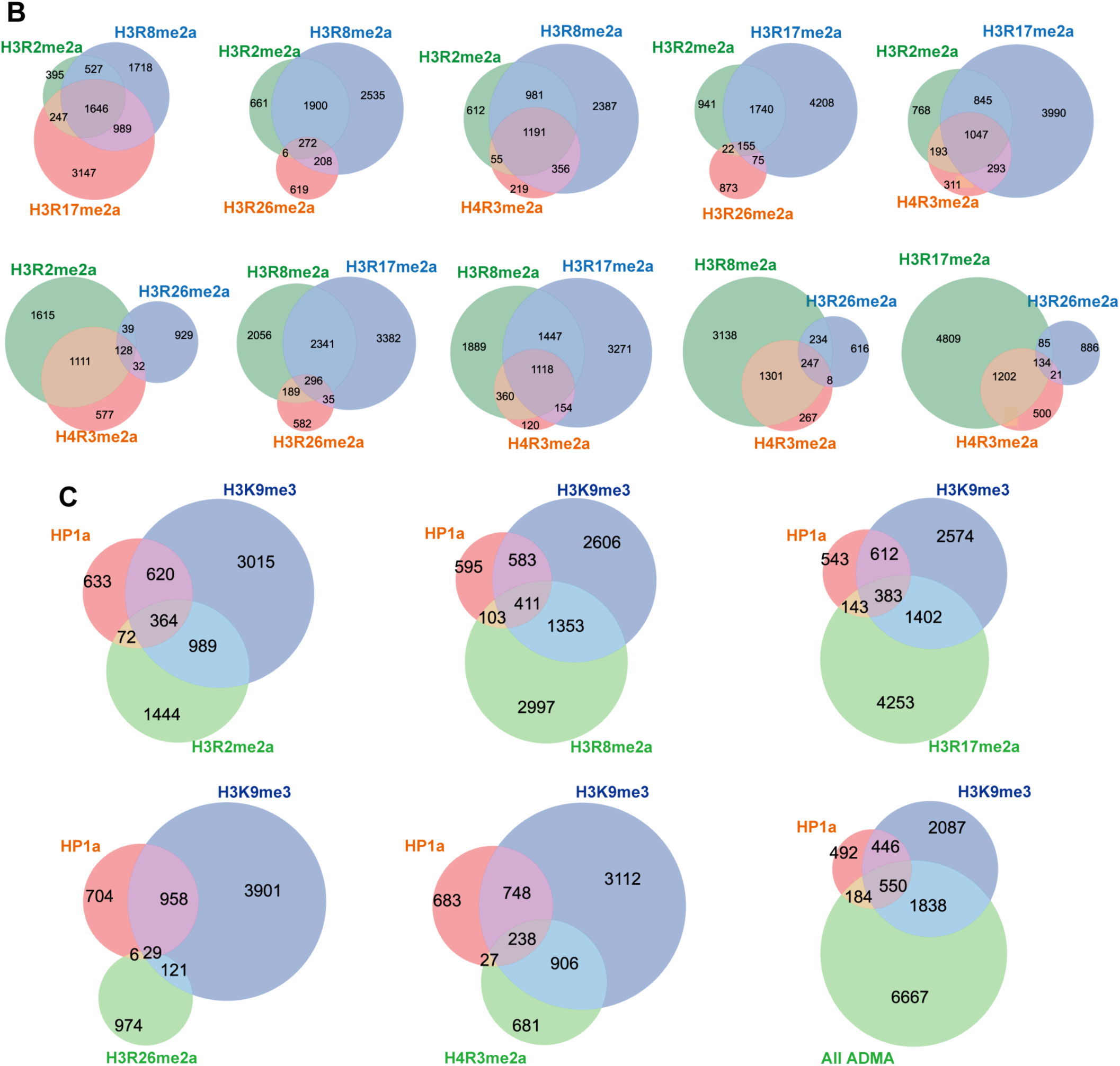
Behavior of ADMA-histones in FLAG-Rhino inducible OSCs. (A) Genome snapshot showing all histone modifications ChIP-seq data in major dual-strand piRNA clusters (42AB, 80F and 102F). The gray-shaded areas indicate the localization sites of FLAG-Rhino in each piRNA cluster in OSCs. (B) Venn diagram showing the sites of overlapping localization of each ADMA-histone. (C) Venn diagram showing the sites of overlapping localization of ADMA-histones, HP1a, and H3K9me3.

**Supplementary Figure 3.**
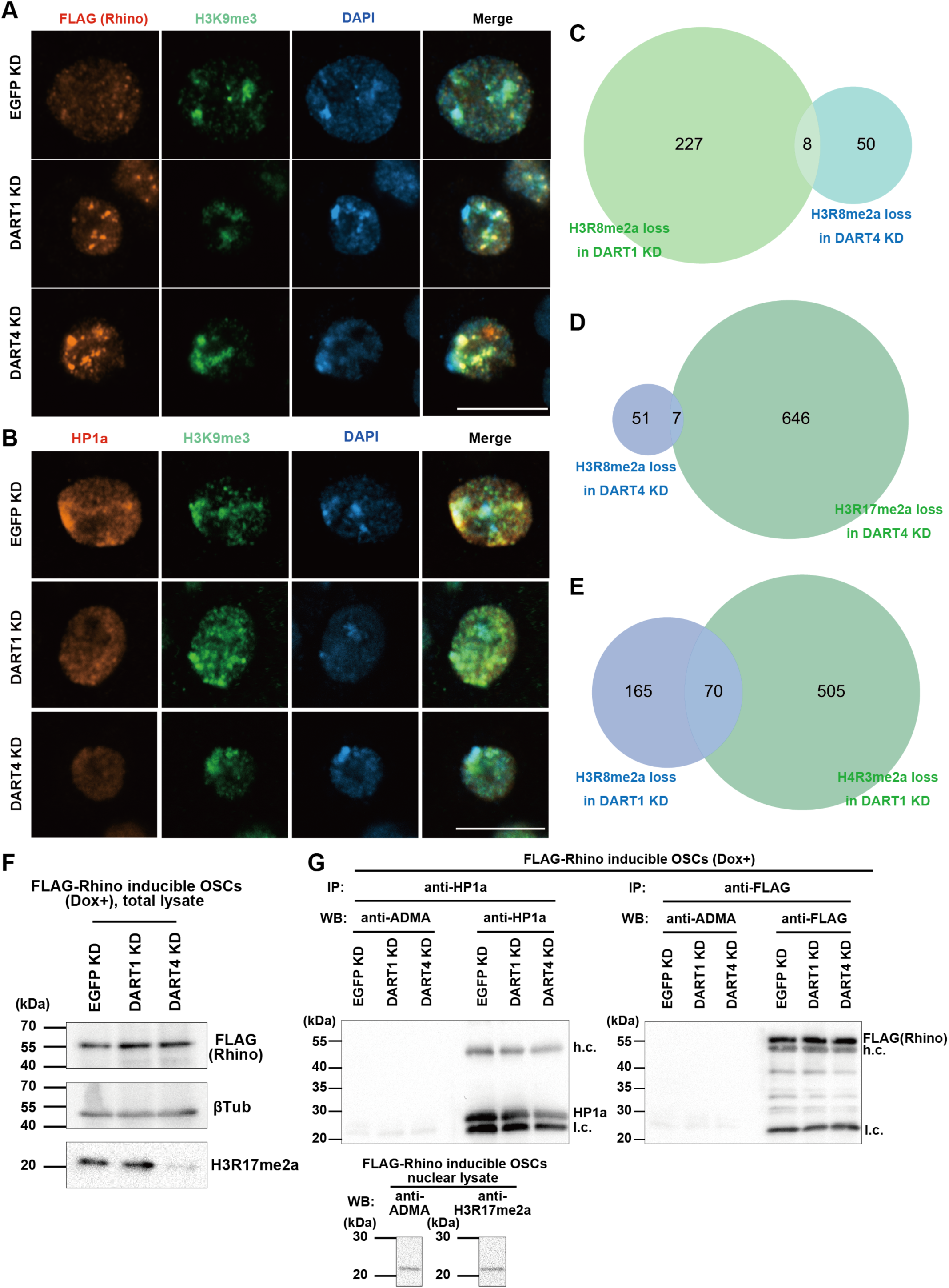
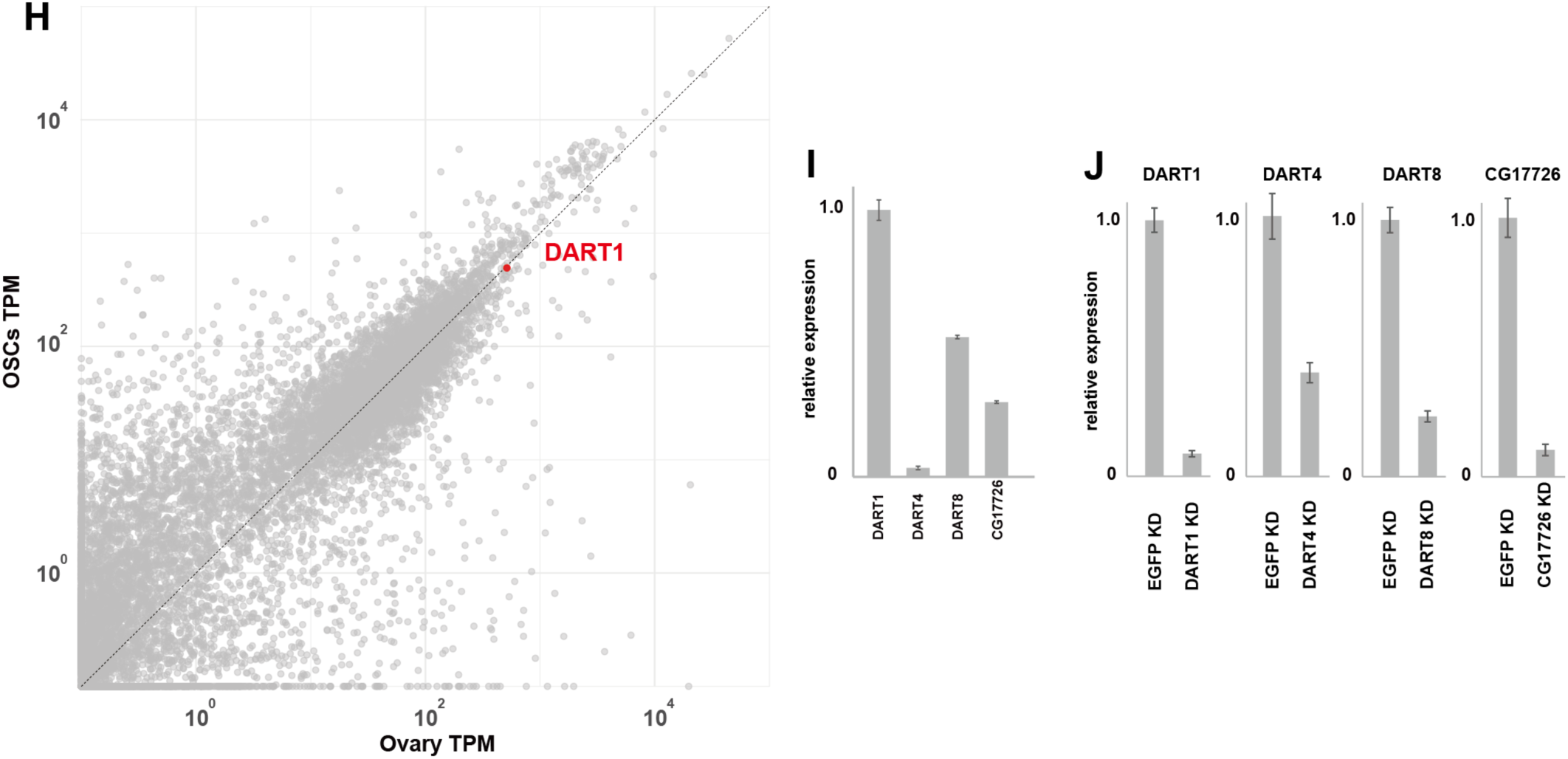
Behavior of FLAG-Rhino, HP1a, and ADMA-histones under DART1-KD and DART4-KD conditions. (A) Confocal immunofluorescent images of FLAG-Rhino (red) and H3K9me3 (green) in FLAG-Rhino inducible OSCs. Dox was added at a final concentration of 5 ng/mL. DAPI shows nuclei (blue). Scale bar represents 10 µm. (B) Confocal immunofluorescent images of HP1a (red) and H3K9me3 (green) in FLAG-Rhino inducible OSCs. Dox was added in final concentration of 5 ng/mL. DAPI shows nuclei (blue). Scale bar represents 10 µm. (C) Venn diagram showing the overlap of the decreased H3R8me2a regions in DART1 KD and DART4 KD. The decreased regions were defined as those with an M value greater than 0.5 in manorm. macs2 callpeak p = 0.01 was used for manorm with input as a control. (D) Venn diagram showing the overlap of the regions where H3R8me2a and H3R17me2a decreased in DART4 KD. The decreased regions were identified using the same conditions as in (c). (E) Venn diagram showing the overlap of the regions where H3R8me2a and H4R3me2a were decreased in DART1 KD. The decreased regions were identified using the same conditions as in (C) and Fig. 3F, G. (F) Western blotting using anti-FLAG, anti-βTub anti-H3R17me2a antibodies on total lysates of FLAG-Rhino inducible OSCs. Dox was added at a final concentration of 5 ng/mL. (G) Western blotting using anti-ADMA, anti-HP1a, anti-FLAG antibodies on immunoprecipitated nuclear lysates of FLAG-Rhino inducible OSCs. Dox was added at a final concentration of 5 ng/mL. Anti-H3R17me2a antibody was used as a positive control for the reactivity of the anti-ADMA antibody. (H) Scatter plot comparing gene expression levels between OSCs and ovaries. Each dot represents a single gene plotted by its TPM values (log10). RNA-seq data for OSCs and ovaries were obtained from DRR05098823 and SRR11879526, respectively. (I) Comparison of qRT–PCR results for DART1, DART4, DART8, and CG17726 in OSCs under EGFP knockdown conditions (n = 3). (J) qRT–PCR analysis of DART1, DART4, DART8, and CG17726 in OSCs under each gene knockdown condition (n = 3).

**Supplementary Figure 4.**
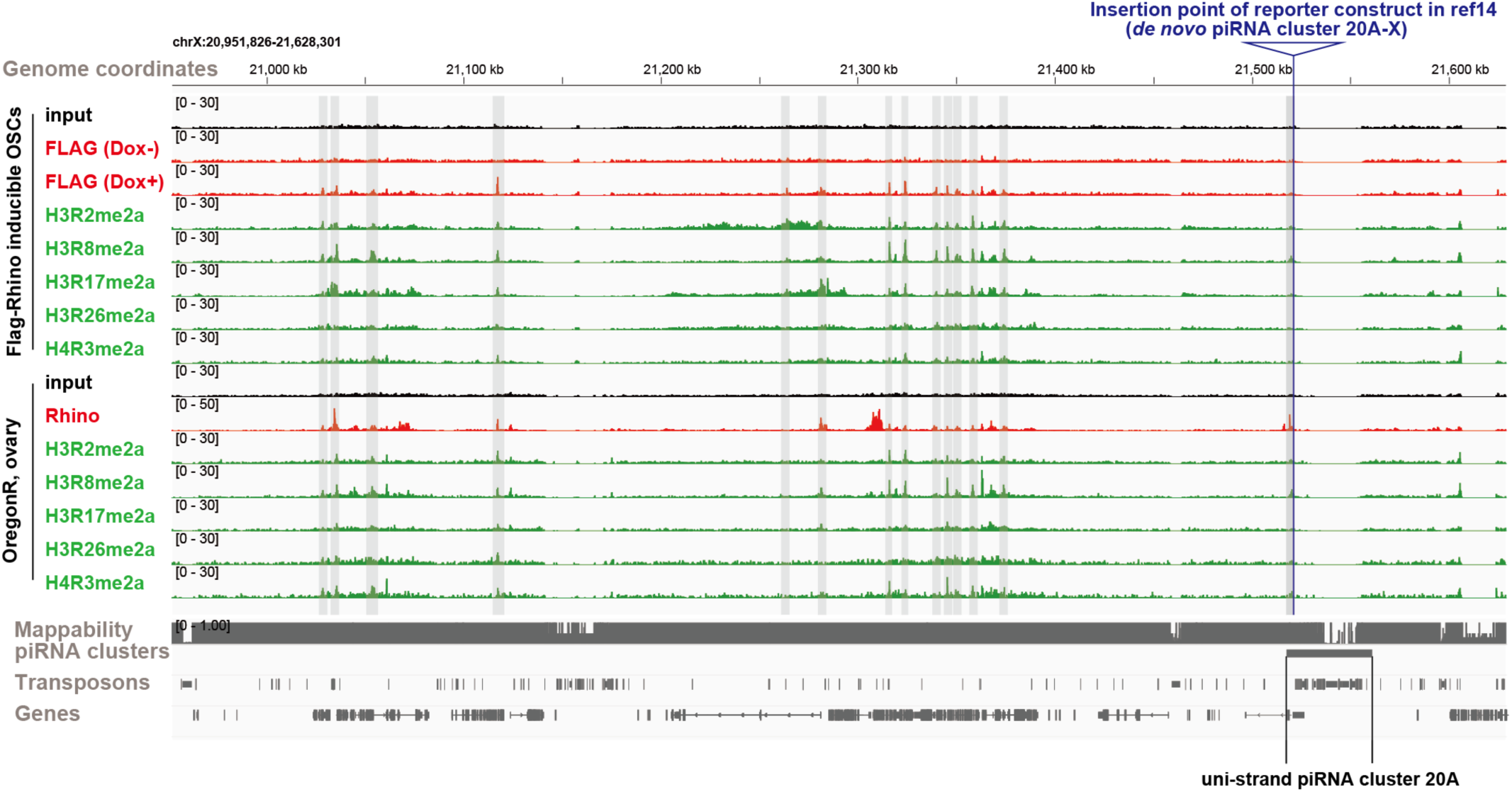
ADMA-histones localize at the ends of Kdm3 GLKD specific ectopic piRNA clusters. Genome browser snapshot showing the localization of H3R2me2a, H3R8me2a, H3R17me2a, H3R26me2a, and H4R3me2a in piRNA clusters in OSCs and Oregon-R ovary. The gray-shaded areas indicate the localization sites of ADMA-histones in Oregon-R ovary.

**Supplementary Figure 5.**
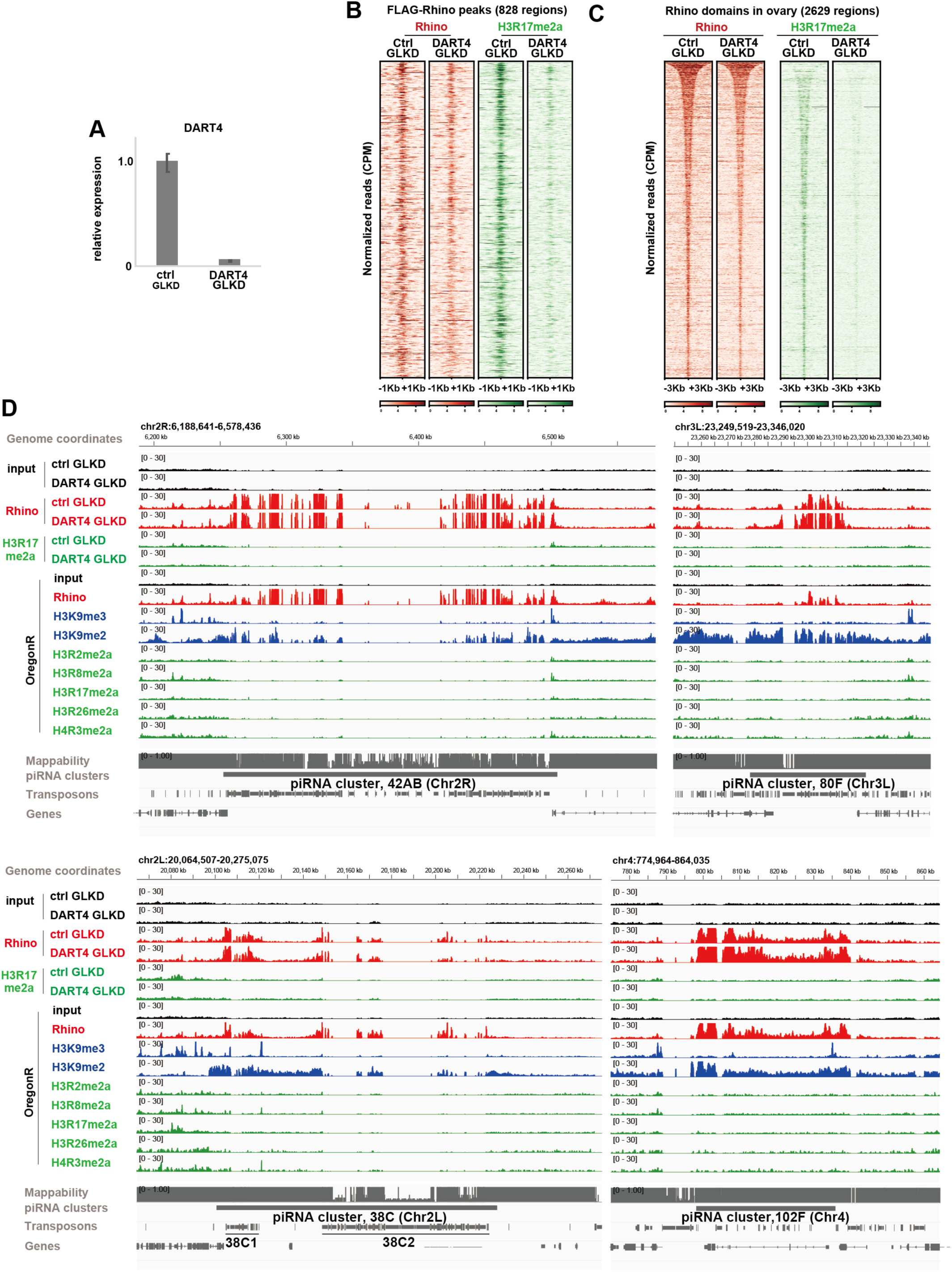
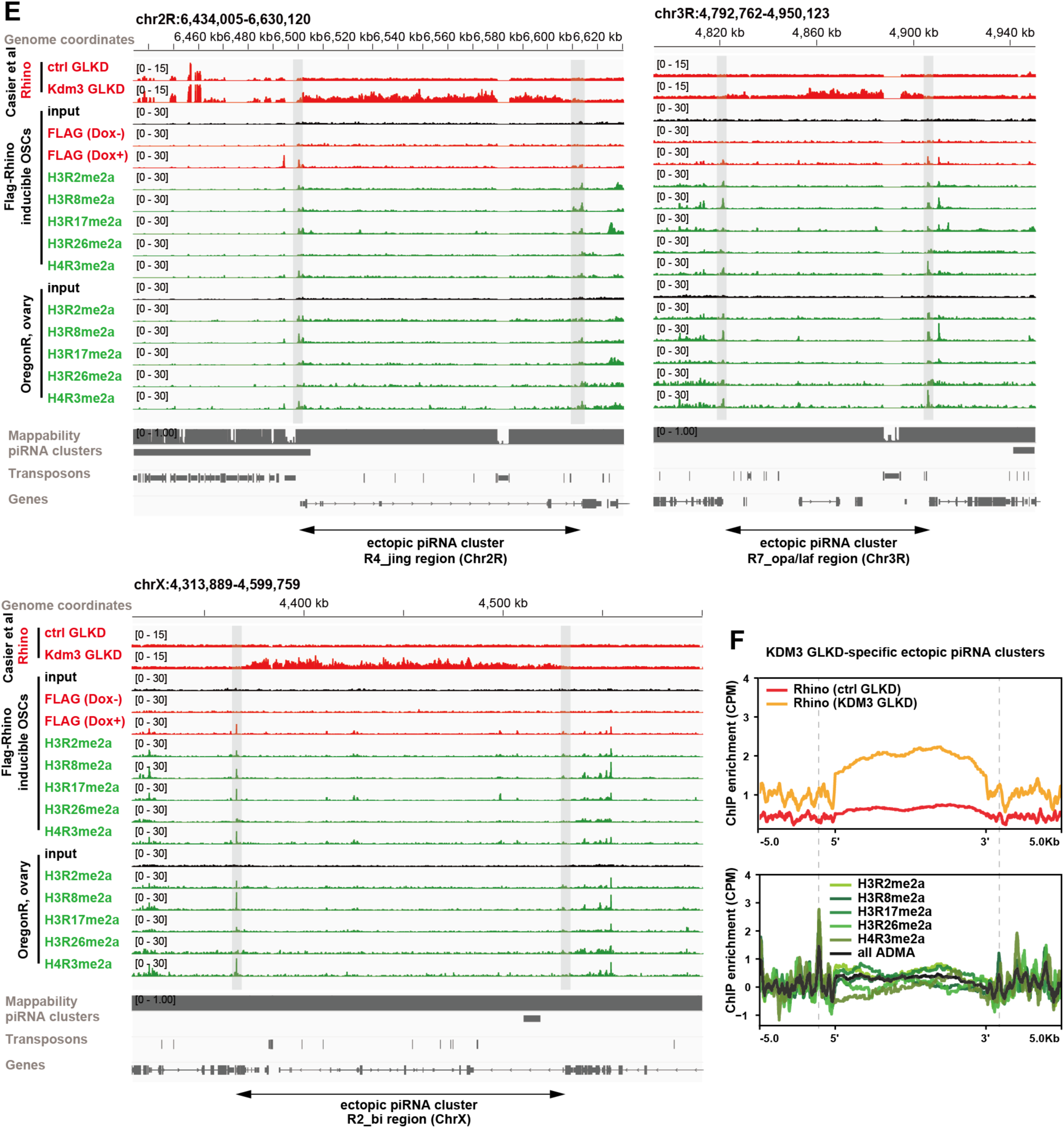
Behavior of Rhino and ADMA-histones under DART4-GLKD conditions. (A) qRT-PCR for DART4 in ovaries under ctrl-GLKD and DART4-GLKD conditions (n = 3). (B) ChIP-seq heatmap showing the localization of Rhino and H3R17me2a in ctrl-GLKD and DART4-GLKD conditions at the FLAG-Rhino localization sites in OSCs. (C) ChIP-seq heatmap showing the localization of Rhino and H3R17me2a in ctrl-GLKD and DART4-GLKD conditions at Rhino localized regions in WT Oregon-R ovary. (D) Genome snapshot of the indicated ChIP-seq data in piRNA clusters, 42AB, 80F, 38C, and 102F. (E) Genome browser snapshot of the indicated ChIP-seq data in representative ectopic piRNA clusters appearing in Kdm3 GLKD (R2_bi region, R4_jing region and R7_opa/laf region). The gray-shaded areas indicate the localization sites of potential FLAG-Rhino considered to be the ends of the ectopic piRNA cluster in OSCs. (F) Metaplot showing ADMA-histones at the major ectopic piRNA clusters R1-R12 (R5 was excluded because of low Rhino localization in Kdm3 GLKD) that appeared in Kdm3 GLKD. Kdm3 GLKD-dependent ectopic piRNA cluster ends were extracted using the following method. Among Rhino peaks listed in the MACS2 callpeak files “GSE203279_MACS2_narrowpeak_Rhino_Ctrl.txt” and “GSE203279_MACS2_ narrowpeak_Rhino_Kdm3GLKD.txt” in the GSE203279 dataset, we first extracted the Rhino peaks that appeared only in the latter^13^. The peaks closest to the 5′ and 3′ ends of the R1-R12 regions in Table S4 of the Casier et al. (2023) were designated as the ends of ectopic piRNA clusters^13^.

**Supplementary Figure 6.**
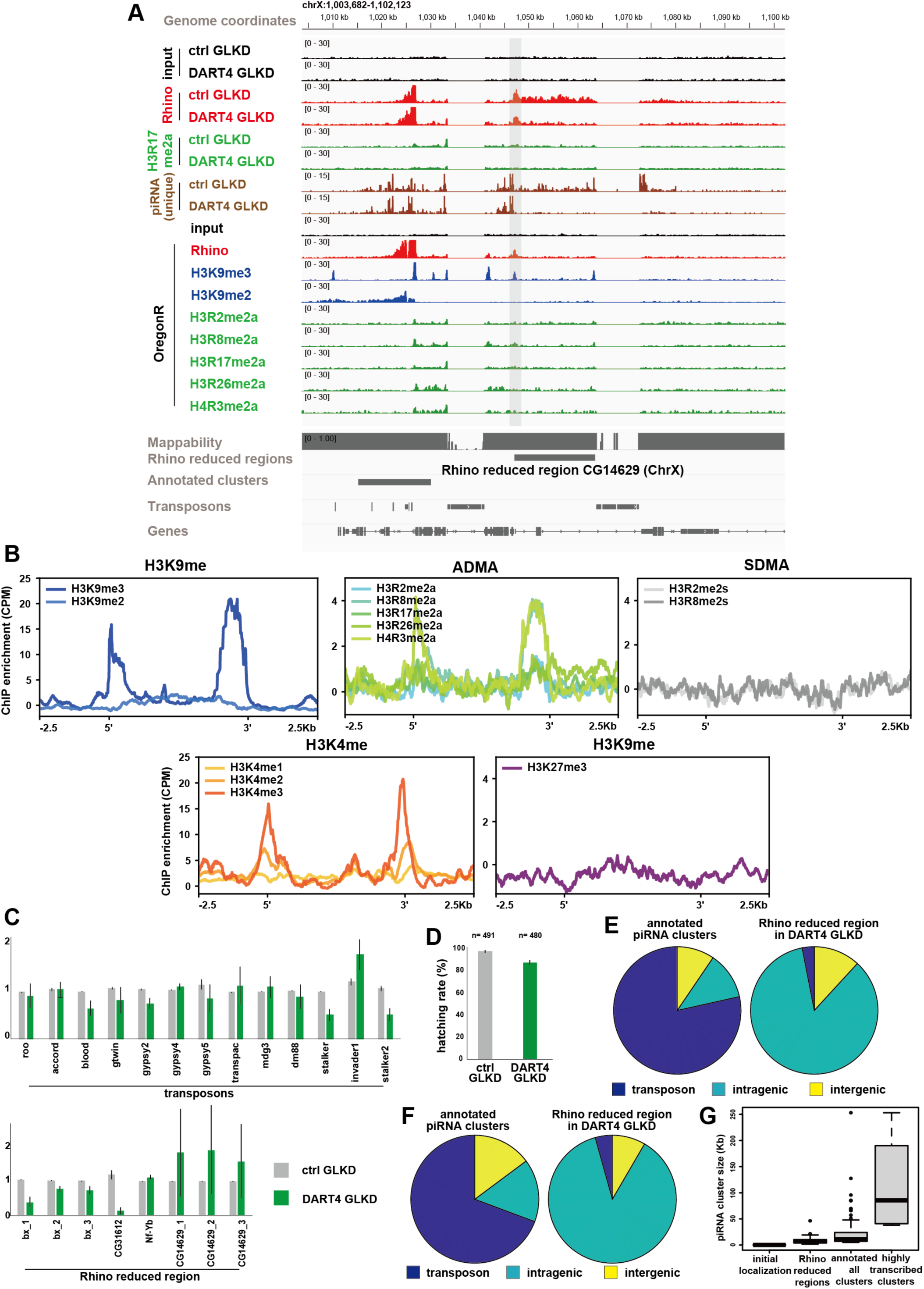
piRNA source loci with H3R17me2a at the ends lose Rhino localization in DART4 GLKD. (A) Genome snapshot of ChIP-seq results of Rhino reduced regions in DART4 GLKD, as shown in Fig. 6D. Rhino reduced region in DART4 GLKD with remarkable loss of Rhino are shown. Black lines indicate boundaries of Rhino reduced regions in DART4 GLKD. The gray-shaded areas indicate the localization sites of potential ADMA-histones considered to be the ends of Rhino reduced regions in DART4 GLKD. (B) ChIP-seq profile showing enrichment of the indicated histone modifications in Rhino reduced regions in DART4 GLKD. ChIP-seq data from WT Oregon-R ovaries were used for each histone modification. SDMA: symmetric dimethylarginine. (C) qRT-PCR showing the expression of transposons and genes within Rhino reduced regions in DART4 GLKD (bx, CG31612, CG14629, and Nf-Yb). Primers are listed in Supplementary Table 1. (D) Egg hatching rate showing fertility in DART4 GLKD. (E) Proportion of transposon/gene sequences contained within Rhino reduced regions in DART4 GLKD and annotated all piRNA clusters. (F) Ratio of piRNA reads mapped to transposons in ctrl-GLKD ovaries^72^ within annotated all piRNA clusters and Rhino reduced regions in DART4 GLKD. The number of piRNA reads in each category is as follows: Transposons in all clusters: 1,020,456; intragenic regions in all clusters: 156,244; intergenic regions in all clusters: 123,370; transposons in Rhino reduced regions in DART4 GLKD: 2,568; intragenic regions in Rhino reduced regions in DART4 GLKD: 71,318; intergenic regions in Rhino reduced regions in DART4 GLKD: 9,907. (G) Box plot showing the lengths of Rhino localization sites in OSCs, Rhino reduced regions in DART4 GLKD, all piRNA clusters annotated in proTRAC (as in Fig. 6B), and highly transcribed piRNA clusters (42AB, 38C, 80F, and 102F).

**Supplementary Figure 7.**
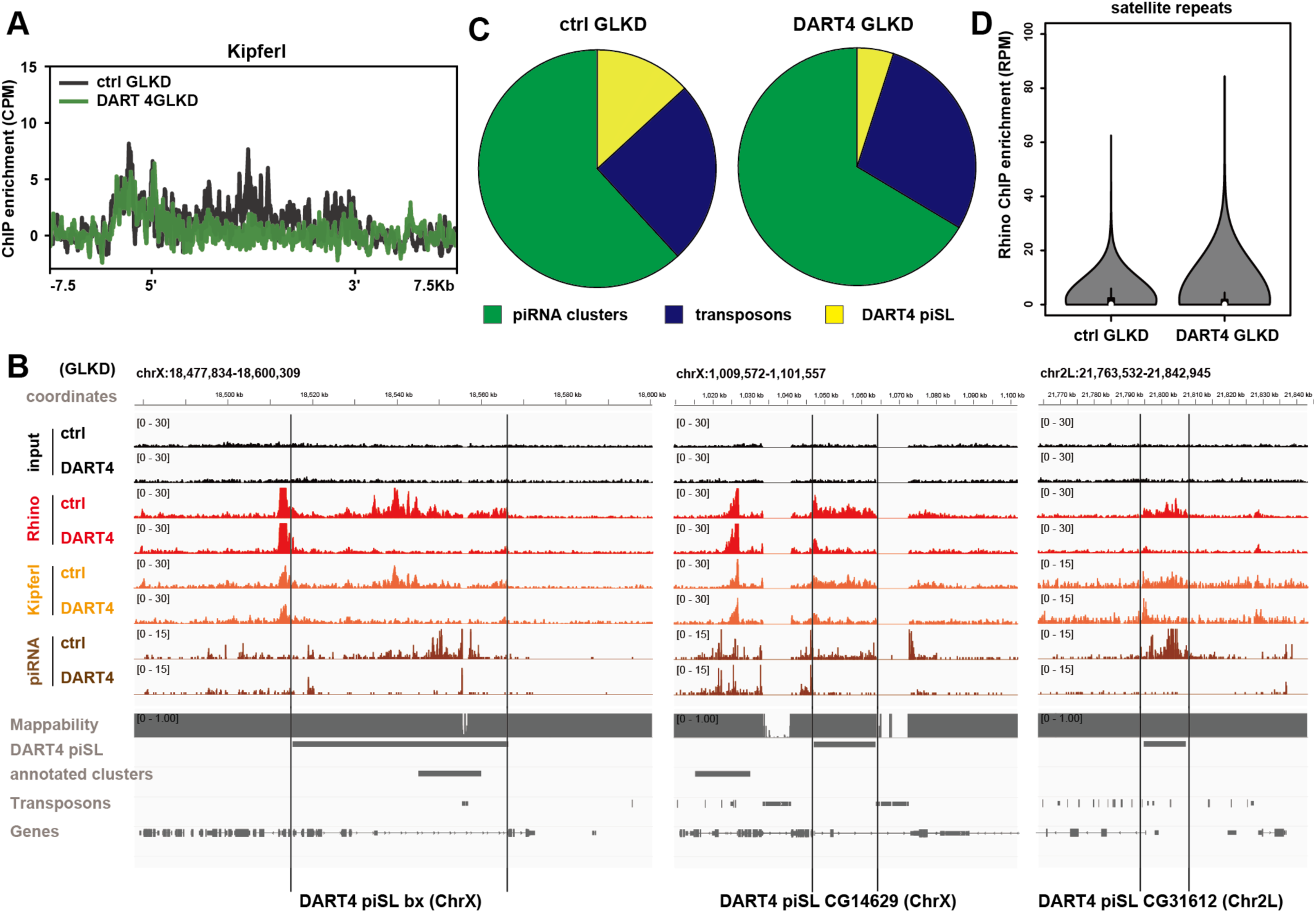
Kipferl and Rhino behavior in DART4 GLKD. (A) ChIP-seq profile of Kipferl enrichment in bx, CG14629, and CG31612 DART4 piSL. (B) Genome snapshot of DART4 piSL in which a significant decrease in Kipferl was observed in DART4 GLKD. (C) Pie chart showing the number of Rhino peaks in piRNA clusters, transposons, and DART4 piSL in ctrl GLKD and DART4 GLKD. The numbers for each are as follows. piRNA clusters in ctrl GLKD: 282; transposons in ctrl GLKD: 114; DART4 piSL in ctrl GLKD: 60; piRNA clusters in DART4 GLKD: 402; transposons in DART4 GLKD: 173; DART4 piSL in DART4 GLKD: 30. (D) Boxplot showing Rhino enrichment in satellite repeats.

**Supplementary Figure 8.**
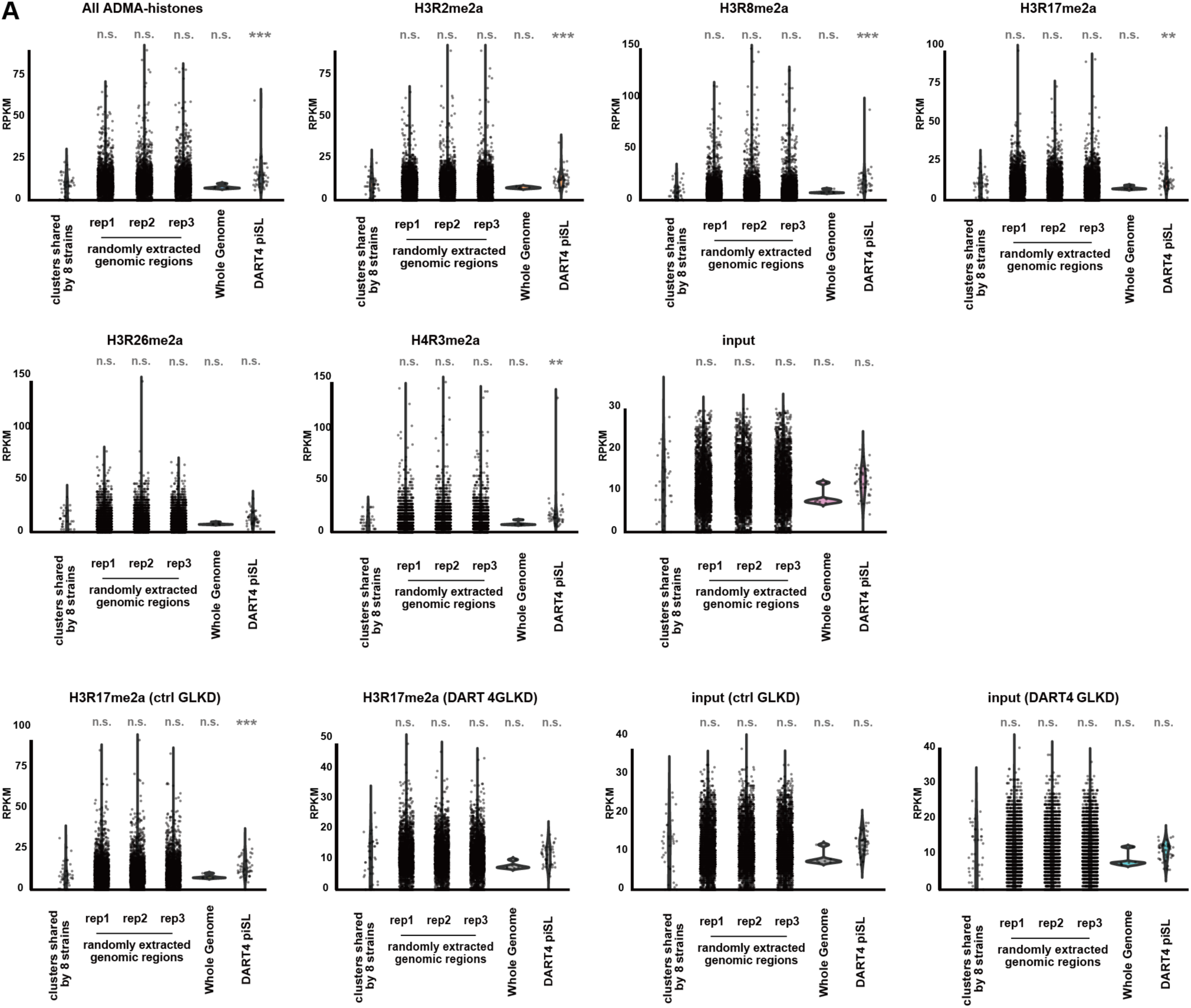
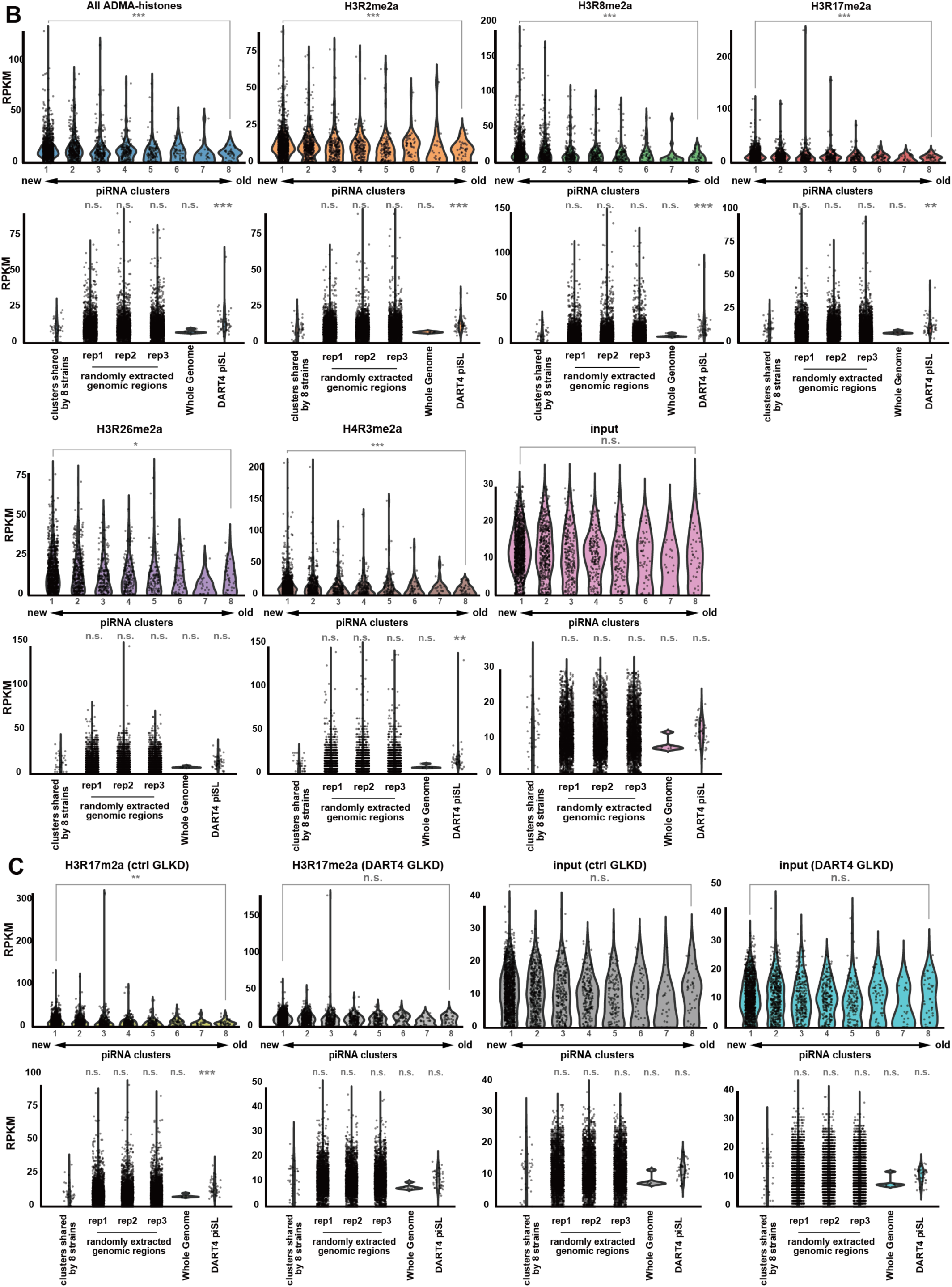
Control for the evolutionary analysis in Fig. 7. (A) RPKM of each ChIP-seq dataset in the piRNA cluster found in all 8 strains (cluster 8, evolutionarily conserved), random genomic regions, the whole genome, and DART4 piSL. Genomic regions with RPKM values above 30 in the input were excluded as they were considered ChIP-seq background noise. For the random genomic regions, 500 bp regions were randomly selected using Python’s random.choice to match the number of piRNA clusters. The whole genome represents the full length of chromosomes 2L, 2R, 3L, 3R, 4, and X. t-test was used for statistical hypothesis testing between clusters found in all 8 strains (cluster 8, evolutionarily conserved) and other indicated genomic regions. Random genomic regions are shown in Supplementary Table 10, and the p-values from the t-test are shown in Supplementary Table 11. (B) Same analysis of Fig. 7A and Supplementary Fig. 8A for each ChIP-seq dataset (Oregon-R) in the piRNA cluster dataset with eight levels of evolutionary novelty in which piRNA reads were detected in Oregon-R strain. (C) Same analysis of Fig. 7A for each ChIP-seq dataset (ctrl GLKD and DART4 GLKD) in the piRNA cluster dataset with eight levels of evolutionary novelty in which piRNA reads were detected in ctrl GLKD strain.

**Supplementary Figure 9.**
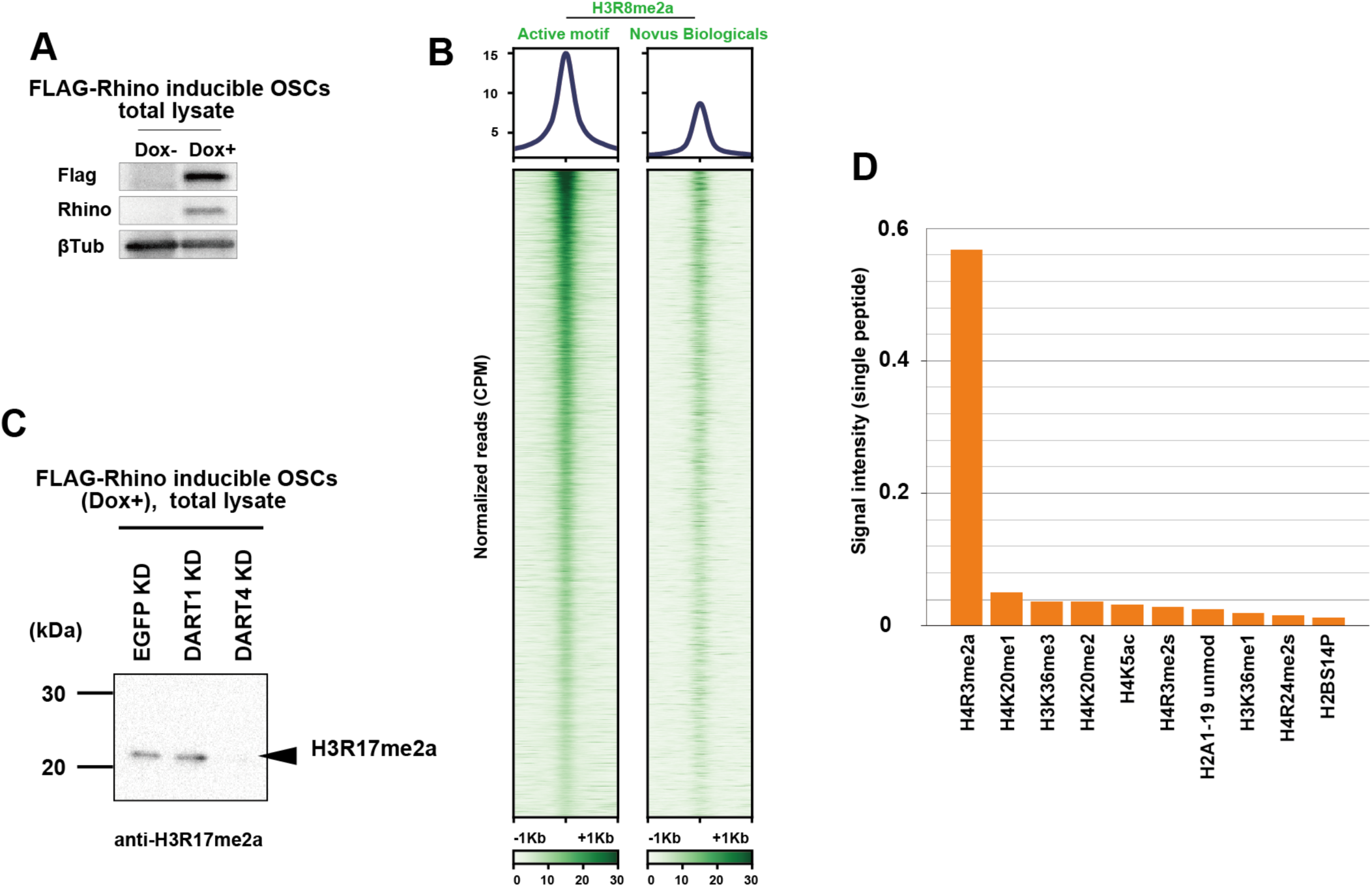
Validation of antibody specificity. (A) Western blotting on total lysates of FLAG-Rhino-inducible OSCs. The anti-Rhino antibody used in immunofluorescence (Fig. 5A) was used. (B) Heatmap showing the localization of H3R8me2a (antibodies produced by different industries) in OSCs at ChIP-seq peaks using the H3R8me2a antibody (Active Motif). The regions of 2 kb around the H3R8me2a localization sites were plotted based on macs2 callpeak results. (C) Western blotting using anti-H3R17me2a antibody on total lysates of FLAG-Rhino inducible OSCs. (D) Histone peptide array using H4R3me2a antibody. The table shows the top 10 single modification peptides with the highest signal intensity.

